# Endocrine therapy induces oxidative stress in ER+ breast cancer that sensitizes persister cells to ferroptosis

**DOI:** 10.1101/2025.07.18.665614

**Authors:** Steven Tau, Malone Friedman, Alyssa M. Roberts, Lauren R. Ferridge, Sierra A. Kleist, Lauren Cressey, Arminja N. Kettenbach, Christopher J. Shoemaker, Todd W. Miller

**Author notes:** Corresponding Author Todd W. Miller, Medical College of Wisconsin 8701 W Watertown Plank Road BSB ROOM 6740, Milwaukee, WI 53226. **Competing interests:** All authors declare no competing interests.

## Abstract

Despite endocrine therapy, recurrence and progression of estrogen receptor alpha (ER)-positive breast cancer remain significant clinical problems. We therefore sought to identify mechanisms underlying endocrine-tolerant persistence. Endocrine-tolerant persister ER+ breast cancer cells were oxidatively stressed during endocrine therapy. Proteomic analysis revealed upregulation of antioxidant-driving enzymes including glutathione peroxidase 4 (GPX4) in persisters. Relief of oxidative stress enhanced persister fitness. The increased oxidative state of persisters enabled lipid peroxidation and ferroptosis. Persisters had an altered lipidome with increased levels of polyunsaturated fatty acids prone to peroxidation, which was attributable in part to increased lysophosphatidylcholine acyltyransferase 3 (LPCAT3, MBOAT5) expression via loss of ER-mediated repression during endocrine therapy. Treatment with the GPX4 inhibitor RSL3 enhanced the anti-persister effects of endocrine-based therapies in xenograft-bearing mice. These findings supporting the development of therapeutic strategies to leverage the oxidative stress induced by endocrine-based therapies and drive ferroptosis as a treatment for ER+ breast cancer.

**Statement of Significance:** Endocrine therapy increases oxidative stress and sensitizes endocrine-tolerant persister ER+ breast cancer cells to ferroptosis, indicating that therapies targeting this metabolic dependency could help prevent disease recurrence and progression.

## Introduction

Tumor formation requires a transformation of cellular metabolism to support heightened proliferation and nutrient demand. Oxidative stress is frequently altered in cancer cells compared to non-transformed counterparts. Oxidative stress is an imbalance of redox status in a cell, namely between reactive oxygen species (ROS) generation and antioxidant scavenging systems. An array factors contribute to high levels of oxidative stress, including higher levels of thymidine phosphorylase (1), NADPH-oxidase activation as a result of RAS oncogene activation, increased proliferative rate, and microenvironmental hypoxia (2). Although cancer cells moderate redox status to survive, ROS can also drive pro-tumorigenic signaling. Thus, cancer cells moderate a balanced level of ROS, often by activation of the transcription factor nuclear factor erythroid 2-related factor 2 (NRF2) to buffer against high levels of oxidative stress (3).

We recently described a reorganization of cellular metabolism that accompanies a switch to a drug-tolerant persister (DTP) state in estrogen receptor alpha (ER)-positive breast cancer (4). Most breast tumors are ER+. Common treatment of early-stage disease involves surgical resection followed by ≥5 yr of adjuvant endocrine therapies that antagonize ER (e.g., tamoxifen) or suppress estrogen biosynthesis (e.g., aromatase inhibitors) to inhibit ER transcriptional activity. However, approximately one-third of patients experience cancer recurrence, many of which occur years after initial diagnosis, suggesting long periods of latency during which DTPs survive despite endocrine therapy. We showed that endocrine-tolerant persister ER+ breast cancer cells have increased mitochondrial content and dependence upon oxidative phosphorylation (4), which has also been observed in other cancer types (5–9). We postulated that a consequence of the increased dependence upon oxidative phosphorylation and decreased glycolysis in endocrine-tolerant persisters may be increased oxidative stress. If so, then oxidative stress may provide a therapeutic vulnerability amenable to pharmacological intervention to help eradicate DTPs and prevent cancer recurrence and progression.

## Materials and Methods

### Cell culture

MCF7, T47D, HCC1428, and ZR75-1 cells were obtained from American Type Culture Collection (ATCC). Cells were maintained in DMEM supplemented with 10% FBS (HyClone Laboratories). Hormone deprivation (HD) involved culture in phenol red-free DMEM containing 10% dextran/charcoal-stripped FBS (DCC-FBS; Hyclone Laboratories) and 2 mM Glutamax (ThermoFisher Scientific). Cell lines were confirmed to be mycoplasma-free (Universal Mycoplasma Detection Kit; ATCC) and authenticated by STR genotyping (University of Vermont Cancer Center DNA Analysis Facility). Cells were treated as indicated with fulvestrant (Selleck Chemicals #S1191), 17β-estradiol (E2; Sigma #E2758), N-acetyl-L-cysteine (Sigma #A9165), reduced L-glutathione (GSH; Sigma #G6013), RSL3 (Cayman Chemical #19288), ferrostatin-1 (Sigma #SML0583), deferoxamine (Sigma #D9533), adrenic acid (Santa Cruz Biotechnology #SC-200784), erucic acid (22:1; Santa Cruz Biotechnology #SC-338740), oleic acid (18:1), PRGL493 (MedChem Express #HY-139180), SC26196 (Cayman Chemical #10792), CP-24879 (Cayman Chemical #10120), BPTES (Cayman Chemical #19284), CB-839 (Cayman Chemical #22038), buthionine sulfoximine (BSO; Cayman Chemical #508228), bafilomycin A1 (Cell Signaling Technology #54645), or tert-butylhydroquinone (tBHQ; Sigma #112941).

### Stable transfection of cell lines

Lentiviral transfer plasmids encoding *LPCAT3* or empty vector control were purchased from Genecopoeia (#EX-Z1415-Lv105-B and # EX-EGFP-Lv181). Lentiviral particles were generated by transfection of Lenti-X cells with transfer plasmid using Lenti-X Packaging Single Shots (Takara Bio). Medium containing viral particles was harvested at 48 h post-transfection and used to infect target cancer cells in the presence of 5 ug/mL polybrene. At 3 d post-infection, stably transfected cells were selected with 1 ug/mL puromycin for 10 d. Cells were also transduced with lentivirus produced from the pHAGE-mt-mKeima transfer plasmid (10) (a gift from Richard Youle; Addgene #131626; RRID:Addgene_131626).

### Fluorescence-activated cell sorting for mtKeima and assay for mitochondrial flux

At 48-72 h after lentiviral infection, cells were sorted with excitation/emission wavelengths of 448/610 nm, and then reseeded in medium containing penicillin/streptomycin. As a negative control, cells were treated with bafilomycin-A1 for 18 h prior to assay. Cells were analyzed by flow cytometry with 405/615 nm and 561/615 excitation/emission wavelengths. Proportions of MitoFlux-positive cells were determined based on 561-nm emission.

### Sulforhodamine B (SRB) assay

This viability assay was performed with modifications from ref. (11). Cells seeded in 96-well plates at 1.5-4e^4^ cells/well were treated as indicated, and drugs/media were refreshed every 3-4 d. At the terminal time point, media were removed, cells were rinsed with PBS, and cells were fixed with 10% trichloroacetic acid at 4°C for 30 min. Wells were then washed with water and allowed to air dry. Wells were stained with 0.04% SRB (Sigma) in 1% acetic acid for 30 min. SRB stain was removed, and wells were washed 5 times with 1% acetic acid and allowed to air dry. SRB was solubilized with 10 mM Tris-HCl pH 7.5, and absorbance at 490 nm was read on a plate reader.

### Serial cell imaging

Cells were seeded in 12– or 24-well plates (1e^3^-1e^4^ cells/well) and treated as indicated. Plates were serially imaged on a Cytation 5 (BioTek). Images were analyzed and cell numbers were quantified with Gen5 software (Biotek).

### Flow cytometry

Cells were plated at 2e^5^ cells/well in 12-well plates. The next day, cells were treated as indicated. Cells were trypsinized and re-suspended in growth medium with either 5 uM CellROX Deep Red (ThermoFisher Scientific #C10422) or 1-2 uM BODIPY 581/591 C11 (ThermoFisher Scientific #D3861). Samples were analyzed either on a MACSQuant-10 (Miltenyi Biotec) or a ZE5 Cell Analyzer (Bio-Rad). Data analysis was performed using FlowJo software.

### Immunoblotting

Cells were lysed in RIPA buffer (20 mM Tris, pH 7.4, 150 mM NaCl, 1% NP-40, 10% glycerol, 1 mM EDTA, 1 mM EGTA, 5 mM NaPPi, 50 mM NaF, 10 mM Na β-glycerophosphate) with HALT protease inhibitor cocktail and 1 mM Na_3_VO_4_. Lysates were sonicated for 10 sec and centrifuged at 17,000 x *g* for 10 min at 4°C. Supernatant was collected, and protein concentration was measured by BCA assay (Pierce). Cell lysates were reduced and denatured using NuPAGE (ThermoFisher Scientific) plus 1.25% β-mercaptoethanol. Fifty ug of protein/sample was analyzed by SDS-PAGE. Protein was transferred onto nitrocellulose membrane and blocked with 5% BSA in TBS containing 0.1% Tween-20 (TBS-T) for 1 h. Membranes were incubated with primary antibody overnight at 4°C on a shaker in blocking solution. Primary antibodies were from Cell Signaling Technology [SLC7A11 (#12691), ER (#8644), GPX4 (#52455), actin (#3700), NRF2 (#12721)] or ThermoFisher Scientific (LPCAT3 #PA-119263). Membranes were then washed with TBS-T 3 times for 10 min, and incubated with HRP-conjugated secondary antibody (Cytiva #NA931V or #NA9340V) in 5% non-fat dry milk in TBS-T for 1 h. After 3 washes with TBS-T for 10 min, signal was developed using ECL Western Blotting Substrate or Supersignal West Pico PLUS substrate (Pierce), and blots were imaged using a Chemidoc MP (Bio-Rad).

### Measurement of GSH/GSSG and NADP/NADPH levels

Cells were seeded in 96-well plates at 0.5-1.5e^4^ cells/well and treated as indicated. Total glutathione and oxidized glutathione were measured separately per manufacturer’s protocol for GSH/GSSG-Glo assay (Promega #V6611). NADP+ and NADPH were measured separately per manufacturer’s protocol for NADP+/NADPH-Glo assay (Promega #G9071). Luminescence was detected on a GloMAX-Multi Detection System (Promega).

### RT-qPCR

RNA was isolated from cells using RNAeasy Universal Plus Mini Kit (Qiagen). One ug of RNA was reverse-transcribed using iScript cDNA Synthesis kit (Bio-Rad #1708890). Real-time qPCR was performed using the iQ SYBR Green SuperMix (Bio-Rad # 1708880) on a Bio-Rad CFX96 thermocycler. Primer sequences were: LPCAT3_fwd: CCAGGAGCTGAGCCTTAACA; LPCAT3_rev: GCAAAGGGGTAACCCAGGAA; ACTB_fwd: TGACAGGATGCAGAAGGAGAT; ACTB_rev: GCGCTCAGGAGGAGCAAT. Data were analyzed using ddCT method.

### Seahorse Assay

Cells were plated at 4-6e^4^ cells/well in Seahorse 96-well plates. The next day, cells were treated as indicated. The Seahorse XF Cell Mito Stress Test kit (Agilent #103015-100) was used per manufacturer’s instructions to calculate basal respiration, maximal respiration, ATP-linked respiration, and spare respiratory capacity. Data were normalized to protein content as measured by BCA assay (Pierce) after cell lysis using RIPA buffer.

### Lipidomics

*Cell treatment*: Cells were treated ± 1 uM fulv for 7 d in 10-cm dishes in triplicate. Medium was then aspirated and cells were washed with PBS followed by 150 mM ammonium acetate. Liquid nitrogen was poured onto cells within each dish to snap-freeze over dry ice, and plates were stored at –80°C. All further steps were performed at University of Michigan Regional Comprehensive Metabolomics Resource Core.

*Internal standards*: The following mass spectrometry-grade lipid standards were used: 1-heptadecanoyl-2-hydroxy-sn-glycero-3-phosphocholine LPC (17:0/0:0), 1,2-diheptadecanoyl-sn-glycero-3-phosphocholine PC (17:0/17:0), 1,2-diheptadecanoyl-sn-glycero-3-phosphoethanolamine PE (17:0/17:0), 1,2-diheptadecanoyl-sn-glycero-3-phospho-L-serine (sodium salt) PS (17:0/17:0), N-heptadecanoyl-D-erythro-sphingosylphosphorylcholine 17:0 SM (d18:1/17:0), cholest-5-en-3ß-yl heptadecanoate 17:0 cholesteryl ester, 1-palmitoyl-2-oleoyl-sn-glycerol 16:0-18:1 DG, 1-heptadecanoyl-rac-glycerol 17:0 MG, 1,2,3-triheptadecanoyl-glycerol triheptadecanoate 17:0TAG, N-heptadecanoyl-D-erythro-sphingosine C17 ceramide (d18:1/17:0), 1,2-diheptadecanoyl-sn-glycero-3-phosphate (sodium salt) 17:0 PA, 1,2-diheptadecanoyl-sn-glycero-3-phospho-(1’-rac-glycerol) (sodium salt) 17:0 PG, 1-heptadecanoyl-2-(5Z,8Z,11Z,14Z-eicosatetraenoyl)-sn-glycero-3-phospho-(1’-myo-inositol) (ammonium salt) 17:0-20:4 PI, 1,3(d5)-dinonadecanoyl-2-hydroxy-glycerol DG d5-(19:0/0:0/19:0) and glyceryl tri(palmitate-d31) TG d31.

*Sample preparation*: Lipids were extracted with a modified Bligh-Dyer method (12) using a 2:2:2 ratio volume of methanol:water:dichloromethane at room temp. after spiking in internal standards. The organic layer was collected and dried under nitrogen. The dried lipid extract was reconstituted in 100 uL Buffer B (10:85:5 acetonitrile/isopropanol/water) containing 10 mM ammonium acetate and subjected to liquid chromatography-mass spectrometry (LC-MS). Internal Standards and Quality Controls: QC samples were prepared by pooling equal volumes of each sample and injected at the beginning and end of each analysis and after every 10 sample injections to provide a measurement of system stability and performance, and of reproducibility of sample preparation method. To monitor instrument performance, 10 uL of a dried matrix-free mixture of the internal standards reconstituted in 100 uL buffer B containing 10 mM ammonium acetate was analyzed. As additional controls to monitor the profiling process, an equimolar mixture of 13 authentic internal standards, a mixture of each experimental sample, and a characterized pool of human plasma were analyzed. Each of these controls were included several times into the randomization scheme.

*Data-dependent LC-MS/MS for measurements of lipids*: Chromatographic separation was performed on a CTO-20A Nexera X2 UHPLC system equipped with a degasser, binary pump, thermostatted autosampler, and column oven (Shimadzu). The column heater temp. was maintained at 55°C and an injection volume of 5 uL was used. For lipid separation, lipid extract was injected onto a 1.8-um particle diameter, 50×2.1-mm id Acquity HSS T3 column (Waters). Elution was performed using acetonitrile:water (40:60) with 10 mM ammonium acetate as Solvent A, and Buffer B with 10 mM ammonium acetate as Solvent B. For chromatographic elution, we used a linear gradient beginning with 60% Solvent A and 40% Solvent B. The gradient was linearly increased to 98% Solvent B over the first 10 min and held at 98% Solvent B for 7 min. Thereafter, solvent composition was returned to 60% Solvent A and 40% Solvent B and held for 3 min. The flow rate used for these experiments was 0.4 mL/min and the injection volume was 5 uL. The column was equilibrated for 3 min before the next injection and run at a flow rate of 0.4 uL/min for a total run time of 20 min.

Mass spectrometry data acquisition for each sample was performed in both positive and negative ionization modes using a TripleTOF 5600 equipped with a DuoSpray ion source (AB Sciex). Column effluent was directed to the ESI source and voltage was set to 5500 V for positive ionization mode and 4500 V for negative ionization mode. The declustering potential (DP) was 60 V and source temperature was 450°C for both modes. The curtain gas flow, nebulizer, and heater gas were set to 30, 40, and 45, respectively (arbitrary units). The instrument was set to perform one TOF MS survey scan (150 ms) and 15 MS/MS scans with a total duty cycle time of 2.4 s. The mass range of both modes was 50-1200 m/z. Acquisition of MS/MS spectra was controlled by the data-dependent acquisition (DDA) function of the Analyst TF software (AB Sciex) with the following parameters: dynamic background subtraction; charge monitoring to exclude multiply charged ions and isotopes; dynamic exclusion of former target ions for 9 s. Collision energy spread (CES) of 20V was set whereby the software calculated the CE value to be applied as a function of m/z. A DuoSpray source coupled with automated calibration system (AB Sciex) was utilized to maintain mass accuracy during data acquisition. Calibrations were performed at the initiation of each new batch or polarity change.

*Data Processing*: Raw data were converted to mgf data format using proteoWizard software (13). The NIST MS PepSearch Program was used to search the converted files against LipidBlast (14,15) libraries in batch mode. We optimized the search parameters using the NIST11 library and LipidBlast libraries and comparing them against our lipid standards. The m/z width was determined by the mass accuracy of internal standards and was set at 0.001 for positive mode and 0.005 for negative mode. The minimum match factor used in the PepSearch Program was 200. The MS/MS identification results from all files were combined to create a library for quantification. All raw data files were searched against this library of identified lipids with mass and retention time using Multiquant 1.1.0.26 (16) (ABsciex). Quantification was done using MS1 data. QC samples were also used to remove technical outliers and lipid species that were detected below the lipid class-based lower limit of quantification. QC samples evenly distributed along analytical runs of the study were analyzed. Missing data were imputed using R package ‘bnstruct.’ Each lipid was normalized using an internal standard that minimized its relative standard deviation (RSD) post-normalization. Each mode (positive and negative) was normalized separately. Normalized data from both modes were combined, and repeats were removed. Data were then normalized to protein content from each sample.

### Animal Studies

Animal studies were approved by the Dartmouth College IACUC (protocol 00002144). Female NOD/SCID/IL2Rγ^-/-^ (NSG) mice were obtained from the Dartmouth Cancer Center Mouse Modeling Shared Resource. Mice were orthotopically injected with 5-10e^6^ MCF7 cells or HCI-003 patient-derived xenograft [PDX; obtained from University of Utah (17)] tumor fragments (∼8-mm^3^) in an inguinal mammary fat pad. An E2 pellet [1 mg in beeswax (18)] was s.c. implanted to induce tumor growth. Tumor dimensions were measured twice weekly using calipers [volume = (length x width^2^/2)]. When a tumor reached 200 mm^3^, mice were randomized to treatment with vehicle, fulvestrant, RSL3, or the combination. Fulvestrant was dissolved in ethanol at 500 mg/mL, diluted in castor oil to a final concentration of 50 mg/mL, and administered s.c. at 5 mg/wk. RSL3 was dissolved in DMSO at 12.5 mg/mL, diluted in PEG-300/Tween-80/saline to a final concentration of 1.25 mg/mL, and administered at 5 mg/kg/d i.p. for 5 d/wk. A subset of mice bearing HCI-003 tumors treated with fulv ± RSL3 were re-randomized to treatment with fulv ± RSL3 ± GDC-0941 (MedChemExpress # HY-50094, 100 mg/kg/d for 5 d/wk, p.o.).

### 13C-glucose tracing metabolic flux analysis

Metabolomics data were previously described (4). Briefly, MCF7 cells were treated with HD medium ± 1 nM E2 for 21 d prior to reseeding in fresh media. Two days later, cells were labeled with media containing U-^13^C-glucose for 45 min. Cells were then washed and snap-frozen, and metabolites were extracted for LC-MS.

### Proteomics

Proteomics data were previously described (4). Briefly, MCF7 cells were treated as indicated, washed with PBS, scraped from dishes and collected by centrifugation, and pelleted cells were snap-frozen. Protein was extracted and trypsin-digested. Peptides were labeled with Tandem-Mass-Tag (TMT) reagent (ThermoFisher Scientific). TMT-labeled peptides were analyzed by LC-MS/MS.

### Statistical Testing

Data were analyzed by t-test (for 2-group experiments) or ANOVA followed by Bonferroni multiple comparison-adjusted post hoc testing between groups (for experiments with ≥3 groups) using Prism software (Graphpad). Tumor growth data were log_2_-transformed prior to linear regression, and growth curve slopes of treatment groups were compared as above.

### Data availability

Lipidomics data are available at the Metabolomics Workbench under accession ST003051. Previously generated Proteomics data are available at ProteomeXchange under accession PXD048700. Previously generated metabolomics data are available at Metabolomics Workbench under accession ST003060. All raw data generated in this study are available upon request from the corresponding author.

## Results

### Endocrine-tolerant persister cells exhibit elevated oxidative stress

We reasoned that increased dependence upon oxidative phosphorylation for ATP generation in ER+ breast cancer cells under selective pressure with endocrine therapy (4) would change the levels of proteins associated with an oxidative stress response (OSR). Hormone deprivation (HD) was used *in vitro* to mimic the estrogen-depleted conditions that occur in patients treated with aromatase inhibitors (19). We leveraged our prior proteomics dataset from MCF7 cells treated with HD ± 1 nM E2 for 4 wk, or with HD for 3 wk followed by E2 for 1 wk (“E2 reversion”). Among a list of 42 genes encoding antioxidant proteins and NRF2/OSR-driven genes (20), our dataset contained 33 proteins. Analysis of the levels of these 33 proteins in MCF7 cells revealed global downregulation in HD compared to E2-replete conditions (ANOVA p<0.0001), suggesting that endocrine-tolerant persisters broadly downregulate antioxidant protein levels (Figure 1A). Furthermore, E2 reversion globally raised the levels of these proteins above those in cells continuously exposed to E2 (p<0.0001), supporting this phenotype as a consequence of persistence during HD.

**Figure 1:**
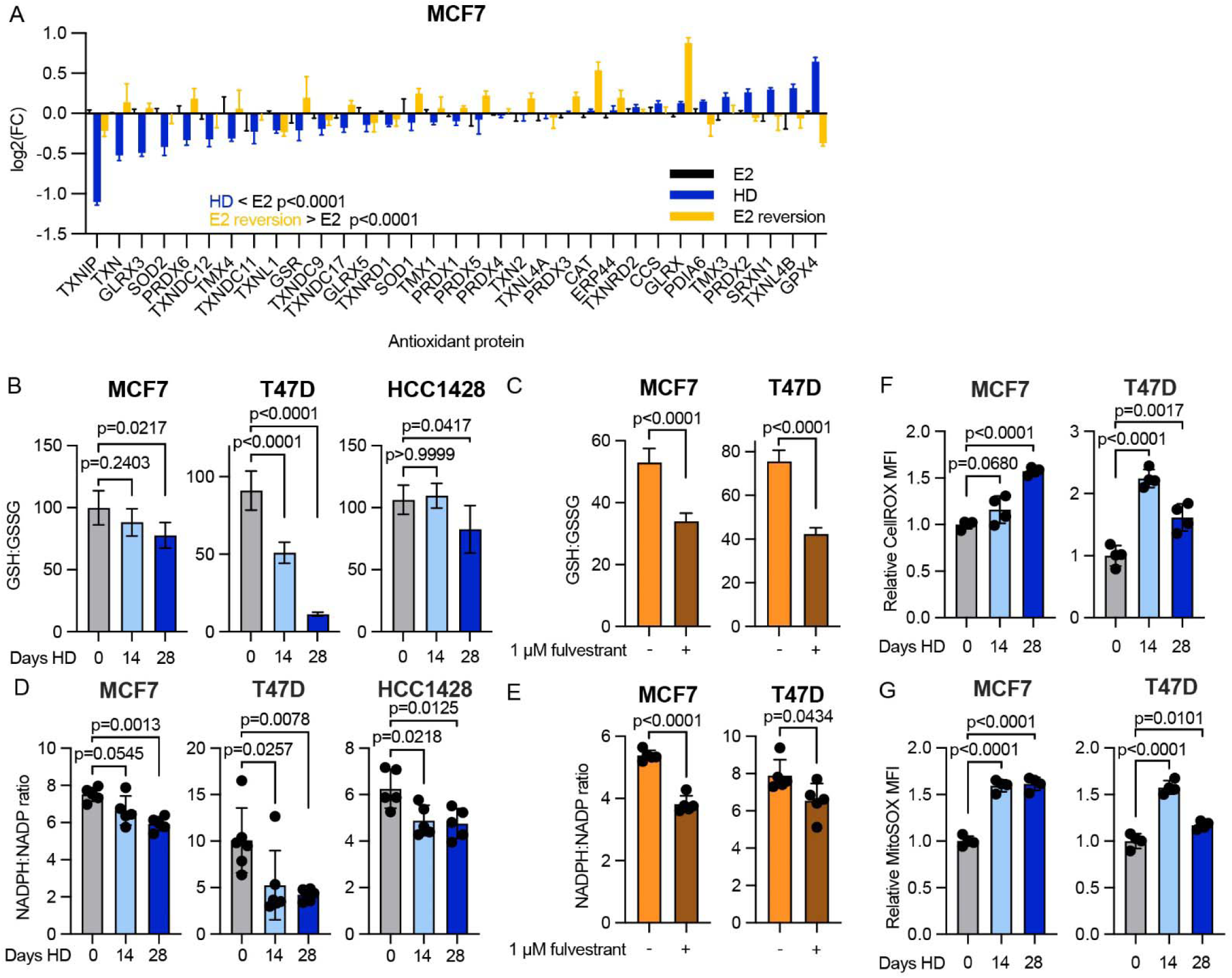
Persister cells surviving endocrine therapy exhibit decreased antioxidant capacity. A) Relative fold change of proteins detected by proteomics assay of cells HD for 4 wk (HD), on estrogen repletion for 4 wk (E2), or reverted with E2 for the last week (E2 reversion). p values were calculated by two-way ANOVA with Bonferroni multiple corrections test (n = 3). B) Ratios of reduced glutathione (GSH) to oxidized glutathione (GSSG) in cell lines pre-HD for 0, 14, or 28 d. C) Ratios of reduced NADPH to oxidized NADP+ in cell lines pre-HD for 0, 14, or 28 d. D) Glutathione ratios in cells pre-treated with fulvestrant for 7 d. E) Ratios of NADP/H in cells pre-treated with fulvestrant for 7 d. F) Relative CellROX median fluorescence intensity (MFI) in cells pre-HD for 0, 14, or 28 d. G) Relative MitoSOX MFI in cells pre-HD for 0, 14, or 28 d. Data are shown as mean + SD in (B/D), n = 5/metabolite.

Glutathione is synthesized by cells to protect against oxidative stress by detoxifying ROS, scavenging oxyradicals, reducing peroxides, and conjugation with electrophilic compounds (21). Through its role in decreasing ROS, the reduced form of glutathione (GSH) becomes oxidized to form glutathione disulfide (GSSG). Replenishment of GSH in cells is dependent upon glutathione reductase and the cofactor nicotinamide adenine dinucleotide phosphate (NADPH). NADPH also has antioxidant properties in scavenging ROS (22). In endocrine-tolerant persisters treated with either HD or the antiestrogen fulvestrant, ratios of reduced to oxidized glutathione were significantly decreased compared to baseline (Figure 1B-C). Additionally, ratios of NADPH (reduced) to NADP+ (oxidized) also decreased in persisters (Figure 1D-E). These data further support an oxidatively stressed state in persisters with shifted antioxidant ratios. Indeed, measurement of relative cellular and mitochondrial ROS levels showed increases in persisters compared to baseline (Figure 1F-G).

In addition to this oxidatively stressed state being driven by ROS levels, we tested for contribution from lower antioxidant levels. We indirectly measured antioxidant production through metabolites traced from [U-^13^C] glucose. Ribose 5-phosphate (R5P) and erythrose 4-phosphate (E4P) are metabolites of the glycolysis-fueled pentose phosphate pathway that aid in the generation of NADPH (22). In line with decreased glycolsis in HD persisters (4), ^13^C-labeled R5P and E4P were also decreased in persisters (Supplemental Figure 1A-B). In addition, persisters showed decreased ^13^C labeling of glutamate, which is converted from the tricarboxylic acid cycle metabolite α-ketoglutarate (Supplemental Figure 1C). Glutamate is crucial for biosynthesis of glutathione both as a precursor and for facilitating cystine import via the cystine/glutamate antiporter xCT (SLC7A11) (23). These data suggest that the underlying cause of oxidative stress in HD persisters is multifactorial: increased production of ROS, including from mitochondria, and decreased antioxidant production.

### Persister fitness is decreased by oxidative state

Since the oxidative state of glutathione was increased in HD persisters (Fig. 1B-C), we evaluated whether glutathione availability perturbs viability. Pre-treatment with HD sensitized cells to pharmacological inhibitors of glutathione synthesis (Figure 2A), including agents targeting glutaminase (GLS1) and glutamate-cysteine ligase (GCL) (Figure 2B). Such agents also increased glutathione oxidation (reflected by the decreased ratios of GSH:GSSG in Figure 2C) and increased ROS (Figure 2D and Supplemental Figure 2A) that trended together.

**Figure 2:**
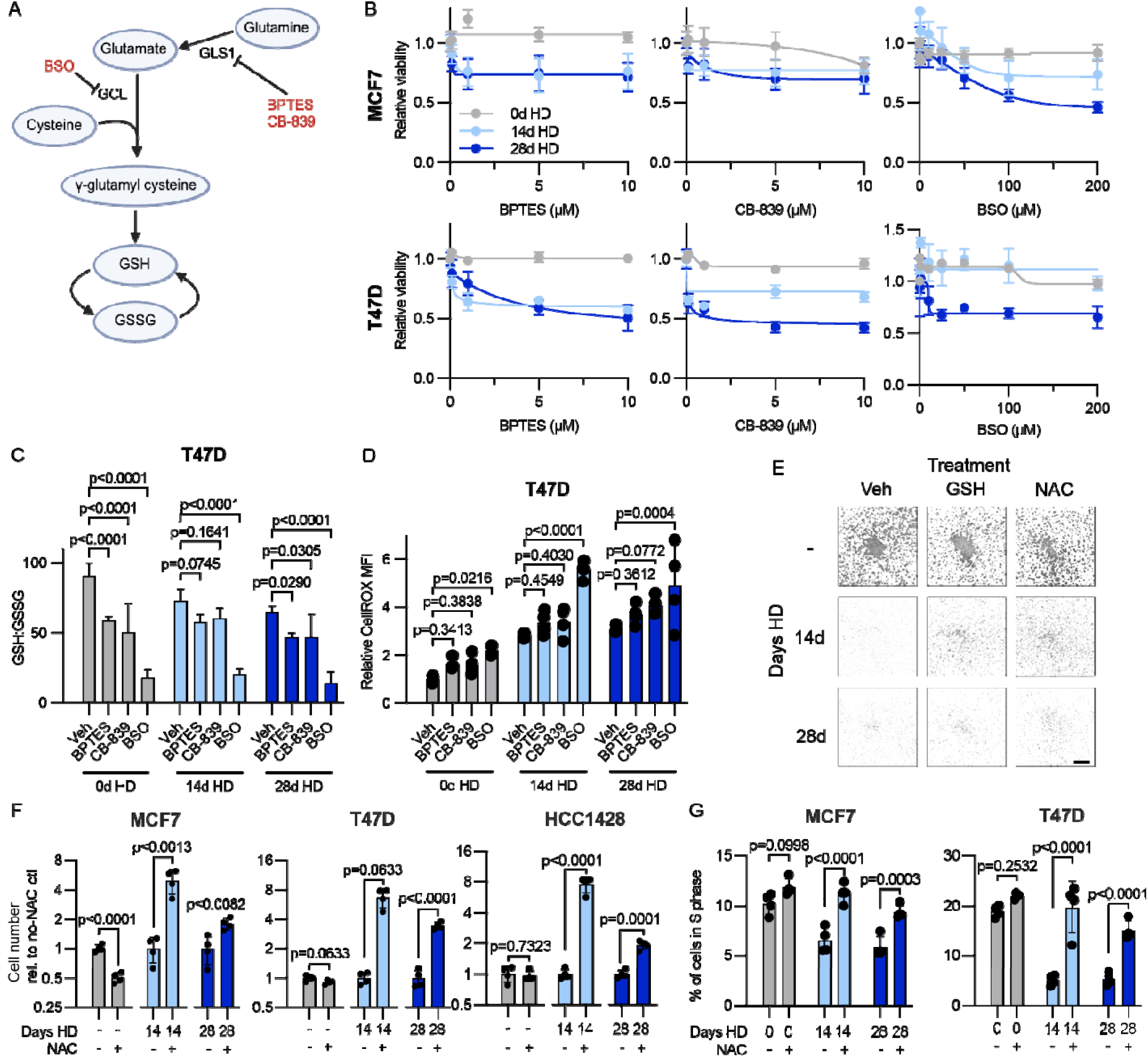
Oxidative state in persisters defines fitness. A) Schematic of glutathione synthesis, with red labels indicating drugs/tool compounds used to inhibit key enzymes. B) Viability curves of MCF7 and T47D cells pre-HD for 0, 14, or 28 d, and subsequently exposed to a range of doses of glutathione synthesis inhibitors. n = 6/dose. C) GSH:GSSG ratios of T47D cells pre-HD for 0, 14, or 28 d and subsequently dosed with 10 μM BPTES, 10 μM CB-839, or 200 μM BSO for 2 d prior to assay. Data represent mean + SD, n = 4/metabolite. D) Relative CellROX MFI of T47D cells pre-HD for 0, 14, or 28 d and subsequently dosed with 10 μM BPTES, 10 μM CB-839, or 200 μM BSO for 2 d prior to assay. E) Representative images of HCC1428 cells pre-HD for 14 or 28 d and treated with 2.5 mM of GSH or NAC for 14 d. As a control, cells kept in estrogen repletion were treated with GSH and NAC for the same time period. Cells were stained with SRB and imaged with Cytation 5. Scale bar = 100 μm. F) Relative numbers of MCF7, T47D, and HCC1428 cells pre-HD for 14 or 28 d and treated with NAC for 14 d. Cell number was quantified by SRB assay. G) Percent of MCF7 and T47D cells in S phase as quantified by cell cycle analysis. Cells were pre-HD for 0, 14, or 28 d subsequently treated with NAC for 3 d prior to processing for cell cycle analysis.

As inhibition of glutathione synthesis decreased growth (Fig. 2B), we tested the effects of the exogenous antioxidants GSH and N-acetyl-cysteine (NAC); NAC can also serve as a precursor for glutathione synthesis (24). Indeed, GSH and NAC each significantly increased HD persister cell number (Figure 2E-F and Supplemental Figure 2B), indicating that buffering of ROS and preventing oxidative stress can partly rescue ER+ BC cells from HD. Interestingly, NAC and GSH also decreased cell fitness in some models under estrogen-replete conditions (Figure 2E-F and Supplemental Figure 2B), supporting the notion that maintaining a homeostatic balance of oxidative state affects viability (25). The NAC-induced increase in persister cell number was paired with an increase in S-phase (Figure 2G), indicating that relief of oxidative stress can induce cell cycling despite withdrawal of estrogen.

We previously reported that HD persisters have increased mitochondrial content (4), which we considered may be associated with oxidative state. Surprisingly, NAC treatment decreased mitochondrial content to near-baseline levels within 3 d in HD persisters (Supplemental Figure 2C). After 3 d of antioxidant treatment, mitochondrial spare respiratory capacity was decreased in HD persisters, but other aspects of respiration were not significantly altered (Supplemental Figure 2D-E). These data suggest (i) the excess mitochondria accumulated in persisters are capable of, but do not contribute to, respiration and (ii) exogenously decreased oxidative stress is associated with decreased mitochondrial content.

Mitochondrial content is dictated by a balance between biogenesis and degradation (collectively termed “mitochondrial dynamics”). The degradation of mitochondria is governed by regulated autophagy (mitophagy) (26). We engineered cells to express mtKeima, a mitophagy reporter protein that accumulates in mitochondria.

The excitation wavelength of mtKeima is shifted when exposed to the acidic pH of a lysosome, allowing for real-time tracking of relative levels of mitophagy (27). Bafilomycin-A1 blocks fusion of autophagic vacuoles with lysosomes (28) and was applied as a control. HD persister cells had greater levels of mitophagy than parental controls (Supplemental Figure 2F), indicating increased mitochondrial dynamics in the persister state.

### Persister fitness is supported by NRF2 activity through oxidative stress response regulation

NRF2 is a master transcriptional regulator of the OSR by driving expression of cytoprotective proteins to reduce ROS (29–32). Kelch-like ECH-associated protein 1 (KEAP1) targets NRF2 for degradation. ROS inhibit KEAP1, enabling NRF2 accumulation (33). We stably knocked down NRF2 in MCF7 cells using shRNA, which was confirmed by loss of NRF2 accumulation upon treatment with the NRF2-activating antioxidant tert-butylhydroquinone (tBHQ) (Supplemental Figure 3A-B). NRF2 knockdown did not impede MCF7 growth rate but decreased persister fitness during HD (Supplemental Figure 3C-D), reinforcing the notion that endocrine therapy creates a persister-specific vulnerability through oxidative stress. NRF2 knockdown increased ROS levels under HD conditions (Supplemental Figure 3E), affirming the role of NRF2 in preventing oxidative stress. Treatment with tBHQ to induce NRF2 increased relative numbers of cells under HD conditions (Supplemental Figure 3F). These data support a role for NRF2 in buffering oxidative stress to enable cell persistence during HD.

### Persister cells are primed for ferroptosis induction

Since persister cells showed increased vulnerability to oxidative stress, we tested the ability of pharmacologic-induced oxidative stress to exacerbate the anti-cancer effects of HD. Glutathione peroxidase 4 (GPX4) was the most strongly upregulated antioxidant-driving protein in HD MCF-7 persisters as observed with proteomics (Fig. 1A). GPX4 catalyzes the reduction of hydroperoxides and lipid peroxides, oxidizing glutathione to prevent oxidative stress. Inhibition of GPX4 can promote accumulation of ROS and iron-dependent lipid peroxidation to trigger ferroptosis, a form of cell death (20,34–36). HD persister cells showed modestly increased sensitivity to the GPX4 inhibitor RSL3 (Supplemental Figure 4A). However, pre-treatment with the antiestrogen fulvestrant, which provides superior inhibition of ER compared to HD (37), increased sensitivity to RSL3 in 3/4 cell lines (Figure 3A). Prior work showed that ferroptosis sensitivity can be dictated by the levels of GPX4 or xCT (SLC7A11) (35). Our analysis showed that fulvestrant-tolerant persisters had decreases in SLC7A11 and/or GPX4 (Figure 3B), possibly contributing to increased sensitivity to RSL3. As with HD, fulvestrant increased ROS levels, and GPX4 inhibition pushed ROS levels higher within 4 h (Figure 3C).

**Figure 3:**
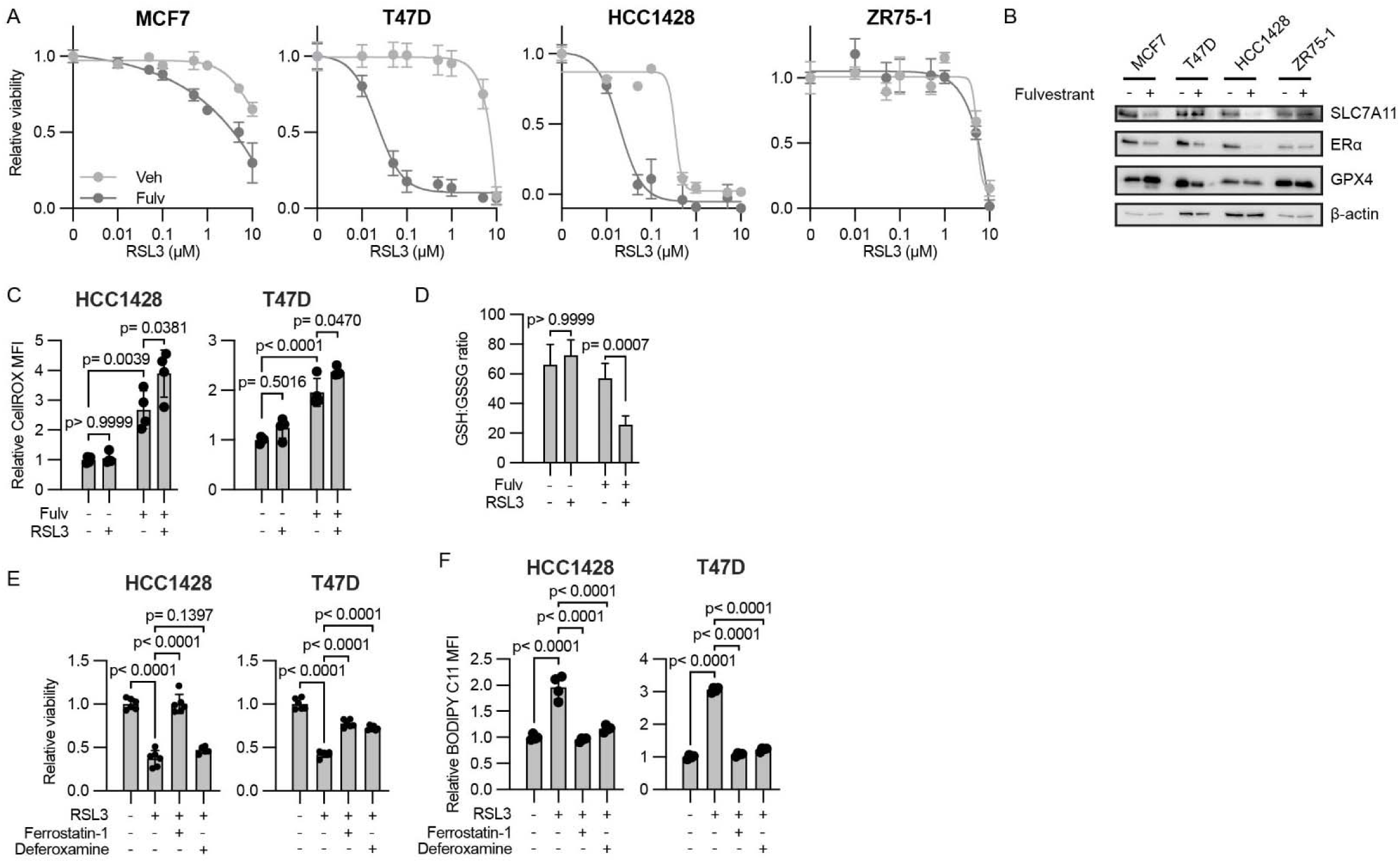
Persister cells are sensitized to ferroptosis induction. A) Viability analysis of cells pre-treated with ± 1 μM fulvestrant before reseeding and treatment with a dose range of RSL3. Cell viability was determined by SRB assay. B) Immunoblot analysis of cells pre-treated with ± 1 μM fulvestrant. C) Flow cytometry analysis of cells pre-treated with ± 1 μM fulvestrant before treatment with ± 1 μM RSL3 for 4 h. Cells were stained and analyzed by flow cytometry for CellROX Deep Red. D) Quantification of GSH:GSSG in T47D cells treated as in (C). Data represent mean + SD, n = 5/metabolite. E) Relative viability of cells persister cells generated by fulvestrant pre-treatment. Cells were reseeded and treated with ± 1 μM RSL3 with ± 10 μM of ferrostatin-1 or deferoxamine. F) Flow cytometry analysis of cells treated as in (E), stained with BODIPY C11, and assayed by flow cytometry.

Similarly, HD increased lipid peroxidation that was driven higher by short-term GPX4 inhibition (Supplemental Figure 4B). Fulvestrant-tolerant persisters showed increased glutathione oxidation that was pushed higher by RSL3 (Figure 3D). These observations collectively suggest that oxidative stress as induced by RSL3 decreases persister fitness.

Lipid peroxidation can be iron-dependent, and iron chelators can abrogate ferroptosis (38). Indeed, the iron chelator deferoxamine and the ferroptosis-preventing antioxidant ferrostatin-1 (39) partly rescued fulvestrant-tolerant persister cells from the growth-suppressive effects of RSL3 (Figure 3E) and prevented RSL3-induced lipid peroxidation (Figure 3F). These data support the sensitivity of endocrine-tolerant persisters to ferroptosis.

## Phospholipid remodeling in persister cells underpins ferroptosis sensitivity

Lipid peroxidation occurs at unsaturated carbon-carbon double bonds, and polyunsaturated fatty acids (PUFAs) are therefore more prone to peroxidation than saturated fatty acids (SFAs) and monounsaturated fatty acids (MUFAs). Not all lipids are peroxidated equally: for example, phosphatidylethanolamines (PEs) with arachidonyl and adrenoyl PUFAs have a higher propensity for peroxidation (40,41). Previous studies correlated ferroptosis sensitivity to the balance of PUFAs vs. MUFAs in lipid membranes (42,43). Given the sensitivity of endocrine-tolerant persister ER+ BC cells to ferroptosis (Fig. 3E-F), we characterized the lipidome to evaluate shifts in lipid composition occurring in persisters. Among 881 lipid species detected in T47D cells, fulvestrant-tolerant persisters showed changes (p≤0.05) in 297 lipid species (Figure 4A). Persisters tended to have shorter fatty acid chain lengths and higher levels of unsaturation (Figure 4B). Focusing on PE FAs as abundant lipids in the plasma membrane, PE PUFAs showed significantly higher levels of most species in fulvestrant-tolerant persisters compared to control-treated cells (Figure 4C). No PE SFAs were significantly altered in persisters compared to controls (Supplemental Figure 5A). PE MUFAs, phosphatidylcholine (PC) SFAs, and PC MUFAs were generally downregulated in persisters, while PC PUFAs showed mixed results (Supplemental Figure 5B-E and Supplemental Table 1).

**Figure 4:**
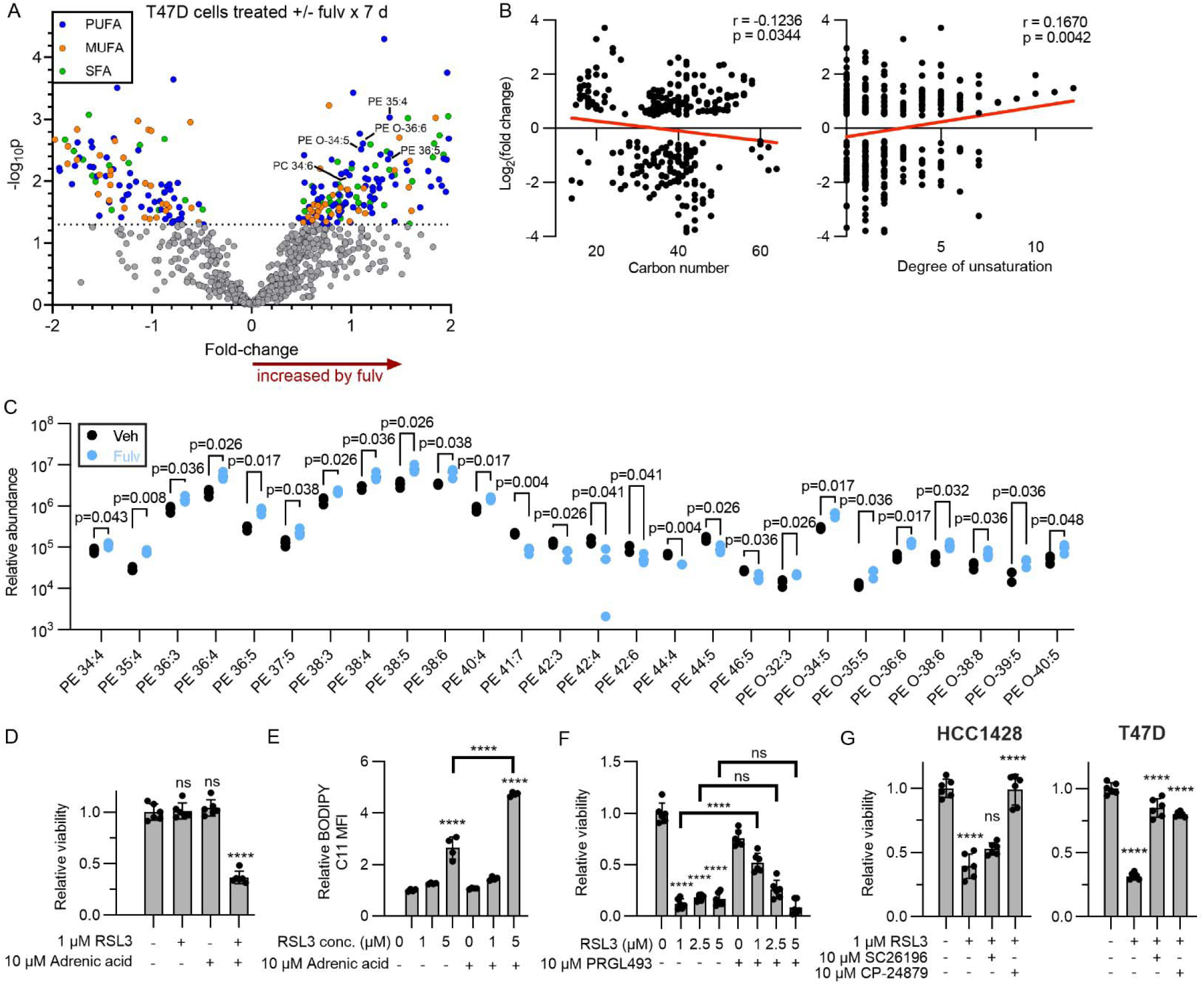
Ferroptosis sensitivity is dependent upon phospholipid content in endocrine-tolerant persisters. A) Volcano plot of lipids differentially abundant in T47D cells treated ± 1 μM fulvestrant for 7 d. Highlighted are lipids with p<0.05. B-C) Pearson correlation plots of lipid carbon number (B) and degree of desaturation (C) with log2(fold change) from (A). Only differential lipids with p<0.05 are included. p is calculated by two-tailed Pearson correlation test. C) Relative abundances of PE PUFA lipid species from (A) that were found with p<0.05 with 5% FDR. D) Relative viability of T47D cells treated with +/− RSL3 and ±adrenic acid for 3 d. Viability was assessed by SRB. E) Flow cytometry analysis of T47D cells treated with RSL3 and adrenic acid for 4 h before staining with BODIPY C11 and flow cytometry. F) Relative viability plot of T47D cells pre-treated with fulvestrant for 7 d before additional treatment with RSL3 and PRGL493. G) Viability plots of cells pre-treated with 1 μM fulvestrant before subsequent treatment +/− RSL3, SC26196, and CP-24879.

We next asked whether lipidome remodeling in persisters influences ferroptosis. Addition of the adrenoyl PUFA adrenic acid to ER+ BC cell culture medium did not affect growth or levels of lipid peroxidation (Figure 4D-E). Although the GPX4 inhibitor RSL3 had no impact on viability, combination with adrenic acid suppressed growth (Figure 4D). In parallel, RSL3 increased lipid peroxidation that was enhanced by adrenic acid (Figure 4E), suggesting that exogenous PUFA was prone to peroxidation that decreased growth.

Acyl-CoA synthetase long chain family member 4 (ACSL4) functionalizes PUFAs through the addition of CoA, enabling inclusion in the cellular phospholipid pool (44). We therefore tested the effects of ACSL4 inhibition with PRGL-493 on viability of fulvestrant-tolerant persister cells. ACSL4 inhibition partially rescued persisters from RSL3-induced growth inhibition (Figure 4F). Delta 5 (Δ5) and Δ6 lipid desaturases are essential for PUFA biosynthesis. We found that inhibition of Δ6 desaturase with SC26196 or Δ5/6 desaturases with CP-24879 partially rescued fulvestrant-tolerant persisters from RSL3-induced growth inhibition (Figure 4G). Since exogenous PUFA increased lipid peroxidation and decreased growth upon GPX4 inhibition (Figure 4D-E), we tested whether exogenous MUFA would be protective. The MUFA oleic acid modestly rescued fulvestrant-tolerant T47D persisters (but not HCC-1428 persisters) from RSL3-induced growth inhibition, but the MUFA erucic acid had no effect (Supplemental Figure 5F). Taken together, these results suggest that lipid unsaturation levels dictate endocrine-tolerant persister ER+ BC cell sensitivity to ferroptosis.

## ER activity regulates LPCAT3 to control PUFA levels

Since PE PUFA levels are upregulated in fulvestrant-tolerant persisters (Figure 4B), we postulated that endocrine therapy alters phospholipid remodeling. Phospholipid composition can be modulated by the selective placement of acyl chains through phospholipid acyltransferases in the Lands Cycle (45). Indeed, membrane-bound glycerophospholipid O-acyltransferases 1 and 2 (MBOAT1/2) have been identified as ER-inducible ferroptosis suppressors by mediating MUFA incorporation into phospholipids. Hence, endocrine therapy can decrease MBOAT1/2 levels, which in turn is expected to increase PUFA levels in the phospholipid pool of ER+ BC cells (43). Investigation of complementary mechanisms of phospholipid remodeling revealed that endocrine therapy increased mRNA and protein levels of LPCAT3 (MBOAT5) in ER+ BC cells (Figure 5A-B). LPCAT3 preferentially binds lyso-PE and lyso-PC to produce membrane phospholipids (46) and favors PUFA-CoAs as acyl donors compared to SFA-CoAs (47). LPCAT3 can thus promote ferroptosis by driving preferential incorporation of PUFA PEs into membranes (48). Indeed, *LPCAT3* mRNA levels were positively correlated with a gene expression signature of ferroptosis across 1,516 cancer cell lines (Supplementary Figure 6A). Mining of MCF7 cistromic profiles of ER and the ER cofactor FOXA1 showed that E2 induces binding of these transcription factors to chromatin regions adjacent to the transcription start site and within the gene body of *LPCAT3*, and E2 suppresses *LPCAT3* mRNA expression (Supplementary Figure 7). In light of our findings with *LPCAT3* mRNA and protein, these data collectively implicate ER in transcriptional repression of *LPCAT3*.

**Figure 5:**
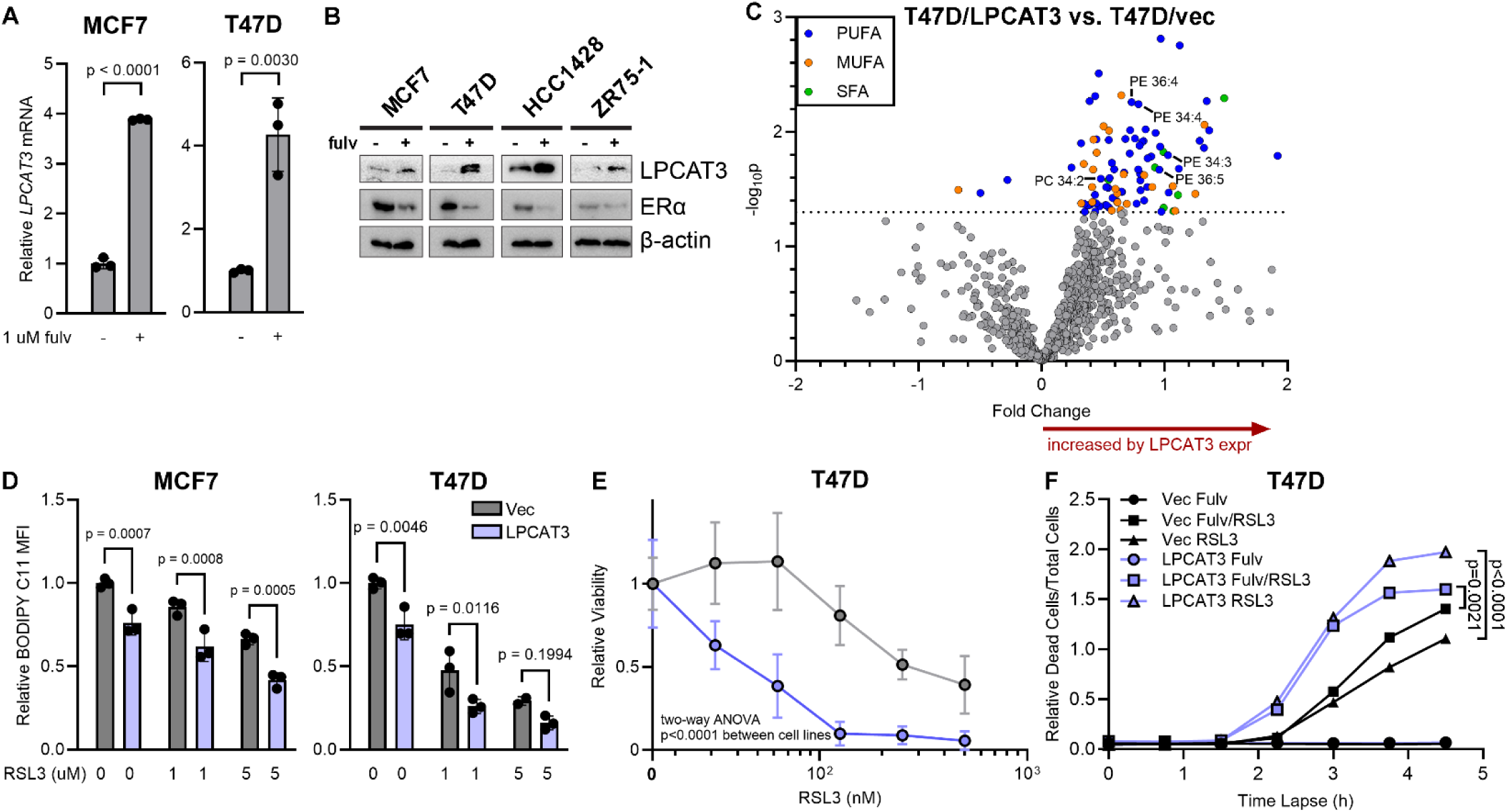
ER signaling regulates LPCAT3 levels that control lipid abundance and cell sensitivity to lipid peroxidation. A) RT-qPCR analysis of cells treated ± 1 uM fulv for 3 d. B) Immunoblot analysis of lysates from cells treated ± 1 uM fulv for 7 d. C) Volcano plot of differentially abundant lipids in T47D cell stably expressing LPCAT3 vs. vector control cells. Highlighted are lipids with p<0.05. D) Cells were treated ± RSL3 for 4 h (MCF-7) or 1 h (T47D), stained with BODIPY C11, and assayed by flow cytometry. E) Cells were treated ± RSL3 for 7 d, and relative growth was measured. F) Cells were pretreated ± 1 uM fulv for 3 d, and then incubated with Cytotox dye and simultaneously co-treated ± 1 uM RSL3. Dye uptake was enumerated as the proportion of total cells. Data in (A/D/E/F) are shown as mean of triplicates ± SD.

We then tested the effects of LPCAT3 expression on the phospholipidome of T47D cells. LPCAT3 overexpression altered (p≤0.05) the levels of 99 of 824 lipid species detected, almost all of which were PUFAs upregulated by LPCAT overexpression compared to vector control (Figure 5C, Supplemental Figure 8, and Supplemental Table 2). In line with the notion that PUFAs are prone to peroxidation, LPCAT3 overexpression increased basal and RSL3-induced lipid peroxidation (Figure 5D), and sensitized cells to RSL3-induced growth inhibition (Figure 5E).

A phenotype associated with ferroptosis is plasma membrane rupture (49). RSL3 treatment induced plasma membrane rupture that started after approximately 2 h and was exacerbated by LPCAT3 overexpression (Figure 5F and Supplemental Figure 9). In a pan-cancer analysis, sensitivity to RNAi-induced silencing of *LPCAT3* was correlated with sensitivity to RSL3 across 235 cell lines (Supplementary Figure 6C). These data suggest that LPCAT3, which is induced by endocrine therapy, increases PUFA content that sensitizes ER+ BC cells to lipid peroxidation and ferroptosis.

### GPX4 inhibition enhances anti-tumor effects of endocrine-based therapy

We extended the evaluation of GPX4 inhibition as a tractable therapy to animal models. Modeling endocrine-sensitive ER+ BC, female mice bearing orthotopic MCF7 tumors were randomized to treatment with vehicle, fulvestrant, RSL3, or both. RSL3 did not significantly alter tumor growth compared to vehicle.

Fulvestrant inhibited MCF7 tumor growth, and combined fulvestrant/RSL3 was significantly more effective than single-agent fulvestrant (Figure 6A and Supplemental Figure 10A). Similarly, combined fulvestrant/RSL3 provided significantly superior efficacy compared to fulvestrant alone in mice bearing T47D tumors (Supplemental Figure 10B).

We then tested GPX4 inhibition in the HCI-003 patient-derived xenograft (PDX) model of ER+ BC. Treatment with fulvestrant or combined fulvestrant/RSL3 significantly slowed tumor growth compared to vehicle, but such growth-inhibitory effects were modest (Figure 6B and Supplemental Figure 10C). We previously reported that HCI-003 tumors had relatively higher activation of the phosphatidylinositol 3-kinase (PI3K) pathway and lower ER levels compared to MCF7 tumors, differences that were exacerbated by fulvestrant treatment. Combined treatment with fulvestrant and the PI3K inhibitor GDC-0941 stabilized HCI-003 tumor volume (50). PI3K pathway-induced mTORC1 activation can prevent ferroptosis via increased MUFA levels (51). We thus postulated that PI3K pathway activation can account for both endocrine and ferroptosis resistance in HCI-003 tumors, and we incorporated GDC-0941 into treatment regimens. Mice treated with fulvestrant ± RSL3 for 28 d (Figure 6B) were randomized to the addition of GDC-0941 (Figure 6C and Supplemental Figure 10D). Treatment with fulvestrant/GDC-0941 significantly inhibited tumor growth compared to single-agent fulvestrant. The triplet of fulvestrant/RSL3/GDC-0941 induced robust tumor regression and was significantly more effective than fulvestrant/GDC-0941 (Figure 6C). These collective observations across tumor models suggest that GPX4 inhibition enhances anti-tumor effects only in the context of other efficacious therapy, supporting a role for GPX4 in persister cells.

## Discussion

Herein, we identified GPX4 as an antioxidant protein upregulated in endocrine-tolerant persister ER+ BC cells compared to control-treated cells. Persisters showed an increased oxidative state that compromised growth. Persisters were sensitized to pharmacological inhibition of GPX4, which increased ROS levels, increased oxidative state, and suppressed growth in an iron-dependent manner. Endocrine therapy via fulvestrant elicited remodeling of the lipidome in persisters that included upregulation of PE PUFAs. Exogenous PUFA enabled growth suppression in the context of GPX4 inhibition that correlated with increased lipid peroxidation; in contrast, inhibition of endogenous PUFA functionalization rescued cells from GPX4 inhibition.

Data mining revealed that endocrine therapy increases LPCAT3 mRNA levels. As a key enzyme in the production of membrane phospholipids, LPCAT3 can drive preferential incorporation of PUFAs. LPCAT3 overexpression induced lipidome remodeling, increasing the levels of lipid species including PUFAs. LPCAT3 overexpression sensitized cells to growth inhibition and ferroptosis induced by GPX4 inhibition. The translational potential of GPX4 as a therapeutic target was confirmed in tumors grown in mice: tumors driven to a persister state were sensitive to GPX4 inhibition.

We examined how ER+ breast cancer cells respond to endocrine therapy as a consequence of induced metabolic reprogramming. We found that persisters remain in a state of oxidative stress marked by a loss in antioxidant stores and higher levels of ROS, coinciding with decreased antioxidant protein levels in persisters (Figure 1). We investigated the consequences of perturbing this balance and found that by instigating more oxidative stress via pharmacological inhibition of glutathione synthesis, persister cell fitness and viability decrease (Figure 2B-D), presumably due to an overwhelming level of oxidation. Contrastingly, amelioration of oxidative stress with exogenous antioxidants improved persister cell fitness and even restarted cell cycling (Figure 3E-F). Other studies have shown that overcoming oxidative stress is a mechanism of recurrence (29). However, the role of the oxidative stress response and signaling pathways such those mediated by NRF2 are complex and contextual (3). Indeed, we saw no or even negative impacts of antioxidants on cells without endocrine therapy (Figure 2F and Supplemental Figure 2E). The underlying mechanisms of how reducing oxidative stress actually improves cell fitness remain incomplete. We uncovered interesting observations on how mitochondria seem to decrease with exogenous antioxidants, and that persister cells have higher levels of mitophagy (Supplemental Figure 2C-F). Further investigation is warranted on how oxidative stress impacts mitophagy. The common understanding is that oxidative stress induces mitophagy to remove damaged mitochondria and decrease ROS (52), which can explain higher levels of mitophagy in persisters, but fail to provide an explanation as to how improving oxidative stress can promote mitochondrial clearance. Furthermore, we demonstrated that the “excess” mitochondria in persisters is not damaged per se and can still respire if forced to (Supplemental Figure 2E-F).

We advanced the investigation ferroptosis induction in endocrine-tolerant persisters and found that persister cells exhibit vastly higher drug sensitivities than parental cells. This phenotype was characterized in other cancer persister models including breast, melanoma, lung, ovarian cancer cells when treated with targeted therapy (20), supporting a generalized phenotype defining persistence. We uncover some underlying differences such as key enzymes involved in ferroptosis such as SLC7A11 and GPX4, and some underinvestigated causes such as lipid composition and its influence in ferroptosis. A shift in lipid levels was detected by lipidomics, with the most pertinent change found in an increase of PUFA PE phospholipids (Figure 4A) that increase the likelihood of lipid peroxidation. Moreover, prevention of PUFA PE buildup, either by ACSL4 inhibition or Δ5/Δ6 desaturase inhibition, can protect against ferroptosis. Interestingly, exogenous MUFAs did not dilute out PUFA PEs in persister cells (Supplemental Figure 5F), suggesting an underlying mechanism that controls this imbalance.

Investigating how the persister state can influence phospholipid composition, we uncovered a regulatory mechanism whereby ER represses the lipid remodeler LPCAT3 to protect against ferroptosis. Furthermore, endocrine therapy inducing the persister state disrupts this repression, causing LPCAT3 levels to increase and presumably incorporate more PUFA PEs into the membrane (Figure 5C), accounting for the increase peroxidation likelihood (48). Finally, we demonstrated ferroptosis induction as a therapeutic approach in combination with fulvestrant in tumors. While MCF7 tumors were moderately responsive to combination treatment, HCI-003 tumors remained insensitive (Figure 6B), presumably due to auxiliary PI3K-AKT-mTOR pathway activation to counter fulvestrant. However, the triplet therapy combination of fulvestrant, GDC-0941, and RSL3 can induce tumor regression in an otherwise treatment-insensitive tumor model (Figure 6C).

It is unknown what the role of ferroptosis suppression is in the context of normal breast biology, and whether estrogen signaling to suppress LPCAT3 levels is required for cells to remain healthy through breast development. Therapeutically, ferroptosis is a promising target based on its selectivity towards persistent cells. Moreover, ferroptosis has been shown to be synergistic with other therapies in breast cancer care, such as radiotherapy by activation of ACSL4 (53). Ferroptosis was demonstrated to be effective in endocrine therapy-resistant models of breast cancer (43), increasing its possible utility to multiple stages of breast cancer progression. However, there is no ideal candidate small molecule ferroptosis inducer designed for translation into the clinic: RSL3 and its analogs bind covalently to GPX4, contributing to its poor pharmacodynamics (54).

One avenue of translating oxidative stress induction with endocrine therapy is to pair it with radiation therapy, known for inducing oxidative stress (55). The evidence surrounding concurrent endocrine therapy and radiation therapy is inconsistent, but hold promise (56). A retrospective study of 41 patients with concurrent endocrine and radiation therapy for breast conservation showed good clinical and pathological responses in the tumors with good cosmetic results (25). However, no trial has been designed to evaluate whether concurrent therapy yields different therapeutic efficacy. Ferroptosis induction, if paired with endocrine and radiation therapy, might yield additional benefit. Furthermore, LPCAT3 levels might be a promising biomarker for ferroptosis sensitivity.

**Figure 6.**
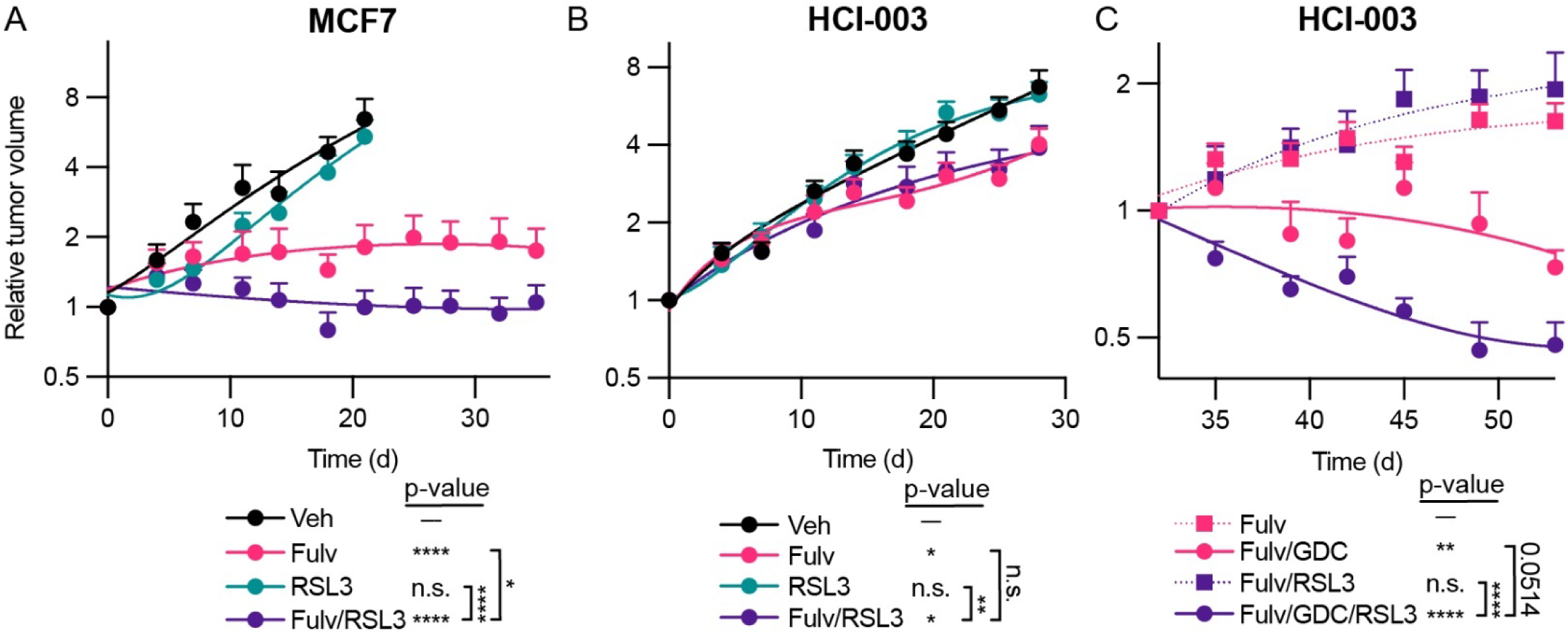
Pharmacological inhibition of GPX4 enhances tumor response to endocrine and PI3K inhibitor therapies. A) Plot of tumor volume curves of MCF7 xenografts treated with ± fulvestrant QW and ± RSL3 QDx5. Data represent mean + SEM (n = 12). B) Plot of tumor volume curves of HCI-003 PDX treated with ± fulvestrant QW and ± RSL3 QDx5. Data represent mean + SEM (n = 10-12). C) Tumor volume plot of HCI-003 xenografts treated with fulvestrant QW ± GDC-0941 QDx5 ± RSL3 QDx5. Data represent mean + SEM (n = 6). Mean starting volume + SD on Day 32 was 345 + 278 mm^3^. Slopes determined by linear regression of log-transformed data were compared by Bonferroni-adjusted posthoc test compared to vehicle unless otherwise indicated with brackets. *p≤0.05, **p≤0.01, ***p≤0.001, ****p≤0.0001. n.s.-not significant.

## Acknowledgements

This work was supported by NIH (R01CA200994, R01CA262232, and R01CA211869 to TWM; Rosalind Borison Memorial Fund and F31CA278418 to ST; R35GM142644 to CJS; Dartmouth College Cancer Center Support Grant P30CA023108; S10OD030242). We thank the following Medical College of Wisconsin Cancer Center Shared Resources for their support: Biorepository & Tissue Analytics; Translational Metabolomics. We thank the following Dartmouth Cancer Center Shared Resources for their support: Mouse Modeling; Pathology (RRID: SCR_023479); Microscopy.

## Figures and legends

**Supplemental Figure 1:**
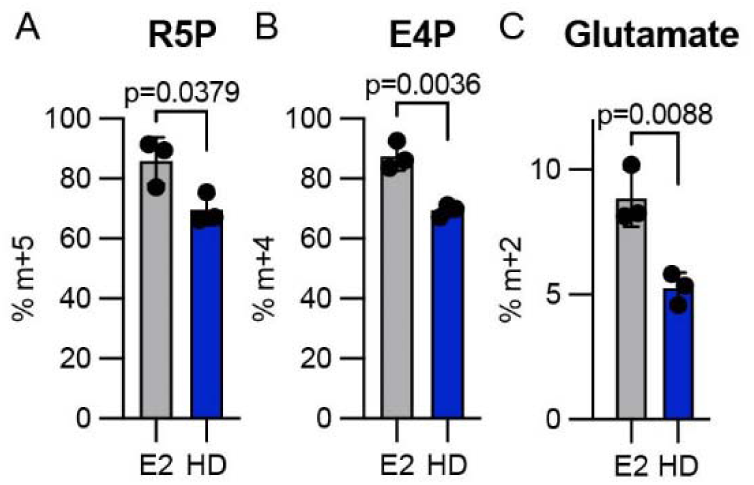
Persisters exhibit decreased pentose phosphate pathway and glutathione precursor metabolites. A-B) Relative labeling of the metabolites ribose 5-phosphate (R5P), erythrose 4-phosphate (E4P), and C) glutamate on samples from treated MCF7 cells as detailed in Chapter 2.

**Supplemental Figure 2:**
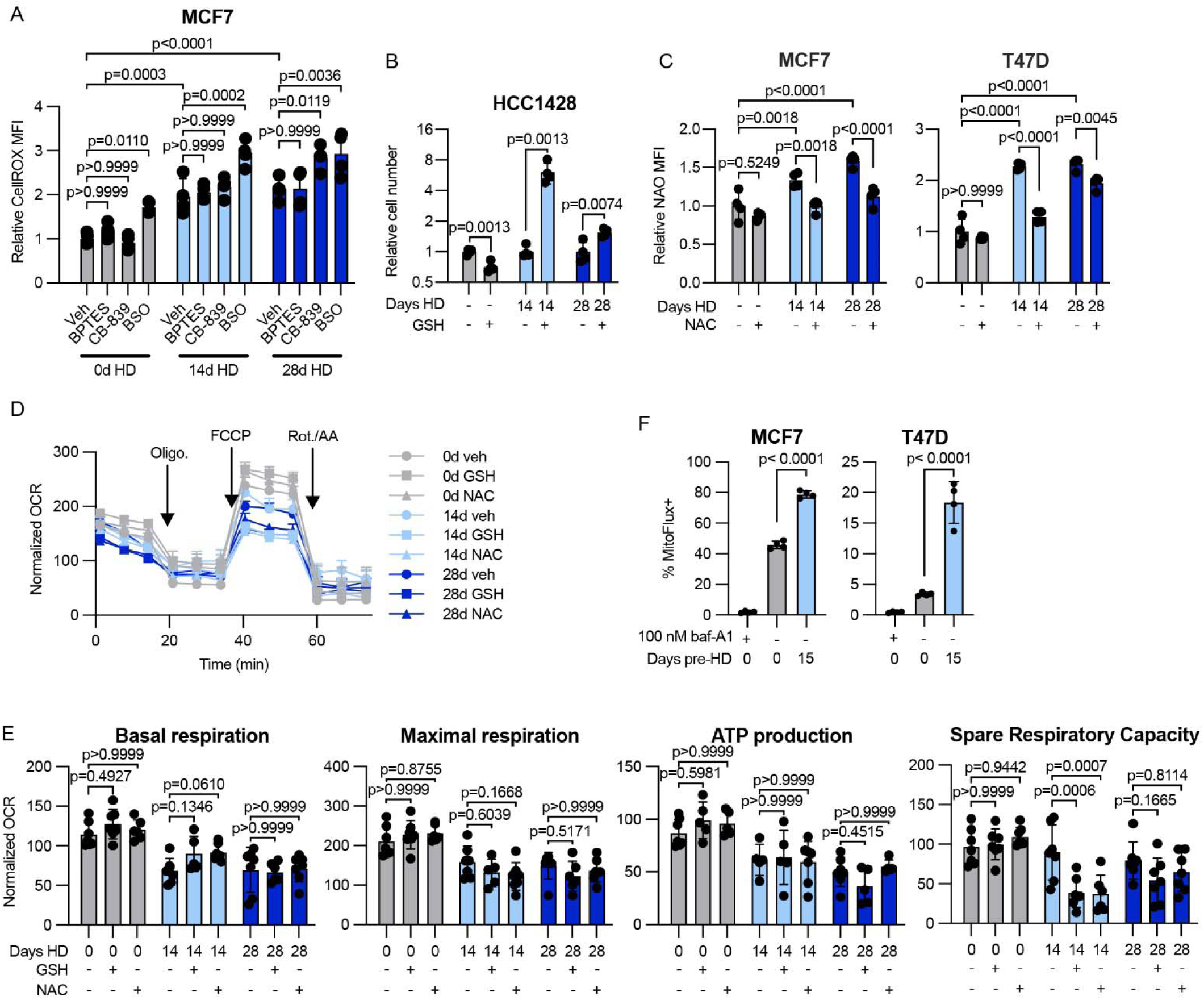
NAC induces removal of excessive mitochondria. A) Relative CellROX MFI of MCF7 cells pre-HD for 0, 14, or 28 d and subsequently dosed with 10 μM BPTES, 10 μM CB-839, or 200 μM BSO for 2 d prior to assay. B) Relative numbers of HCC1428 cells pre-HD for 14 or 28 d and subsequently treated with GSH for 14 d. As a control, HCC1428 cells were treated with GSH in hormone repletion. C) Relative NAO MFI of MCF7 and T47D cells pre-HD for 0-28 d and subsequently treated with 5 mM NAC for 3 d. Cells were stained with NAO and assayed by flow cytometry. D-E) Seahorse analysis of HCC1428 cells pre-HD for 0-28 d prior to treatment with 5 mM GSH or NAC. Mitochondrial respiration was assessed by Seahorse for OCR using the Mito Stress Test Kit. Data represent mean + SEM with *n* = 6/treatment group. F) Flow cytometry analysis of MCF7 and T47D mtKeima cells. Cells were engineered to express mtKeima and pre-HD for 0-15 d. Cells were plated and subsequently treated with baf-A1 for 18 h prior to assay. Cells were analyzed by flow cytometry, and baf-A1-treated cells were used as a negative control to identify MitoFlux+ cells.

**Supplemental Figure 3:**
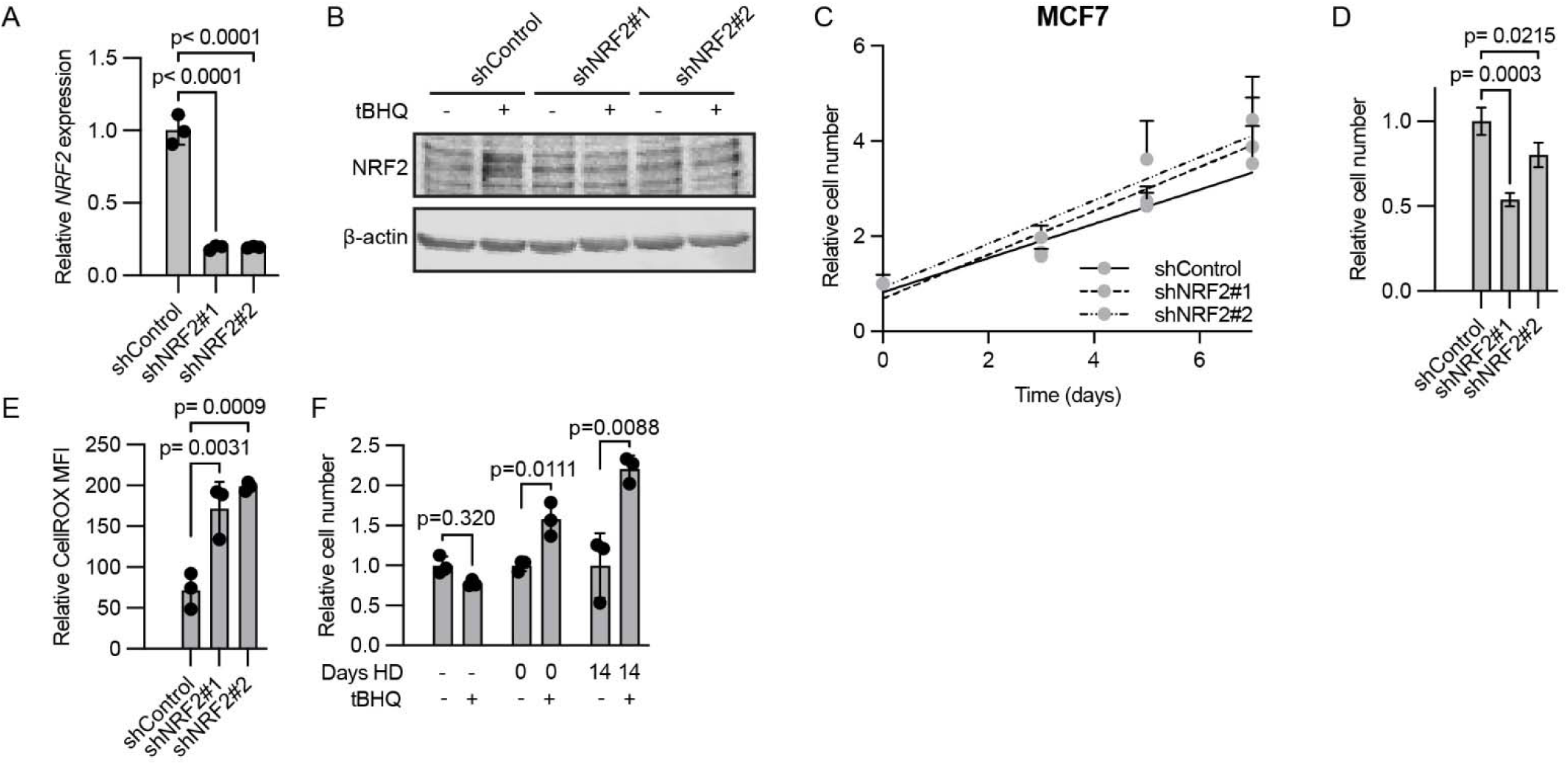
NRF2 activation increases persister cell fitness. A) qPCR quantification of MCF7 cells with NRF2 knockdown. B) Immunoblot of MCF7 cells with NRF2 knockdown and treated with 20 μM tBHQ for 2 d. C) Quantification of MCF7 relative cell number in growth media over 7 d. n = 6. D) Quantification of MCF7 relative cell number in HD for 14 d. n = 6. E) Flow cytometry analysis of cells pre-HD for 14 d prior to staining and flow cytometry with CellROX Deep Red dye. F) Quantification of relative number of cells under estrogen repletion (−) or pre-HD for 0 or 14 d, and then treated +/− 20 μM tBHQ for 14 d with the same hormone conditions.

**Supplemental Figure 4:**
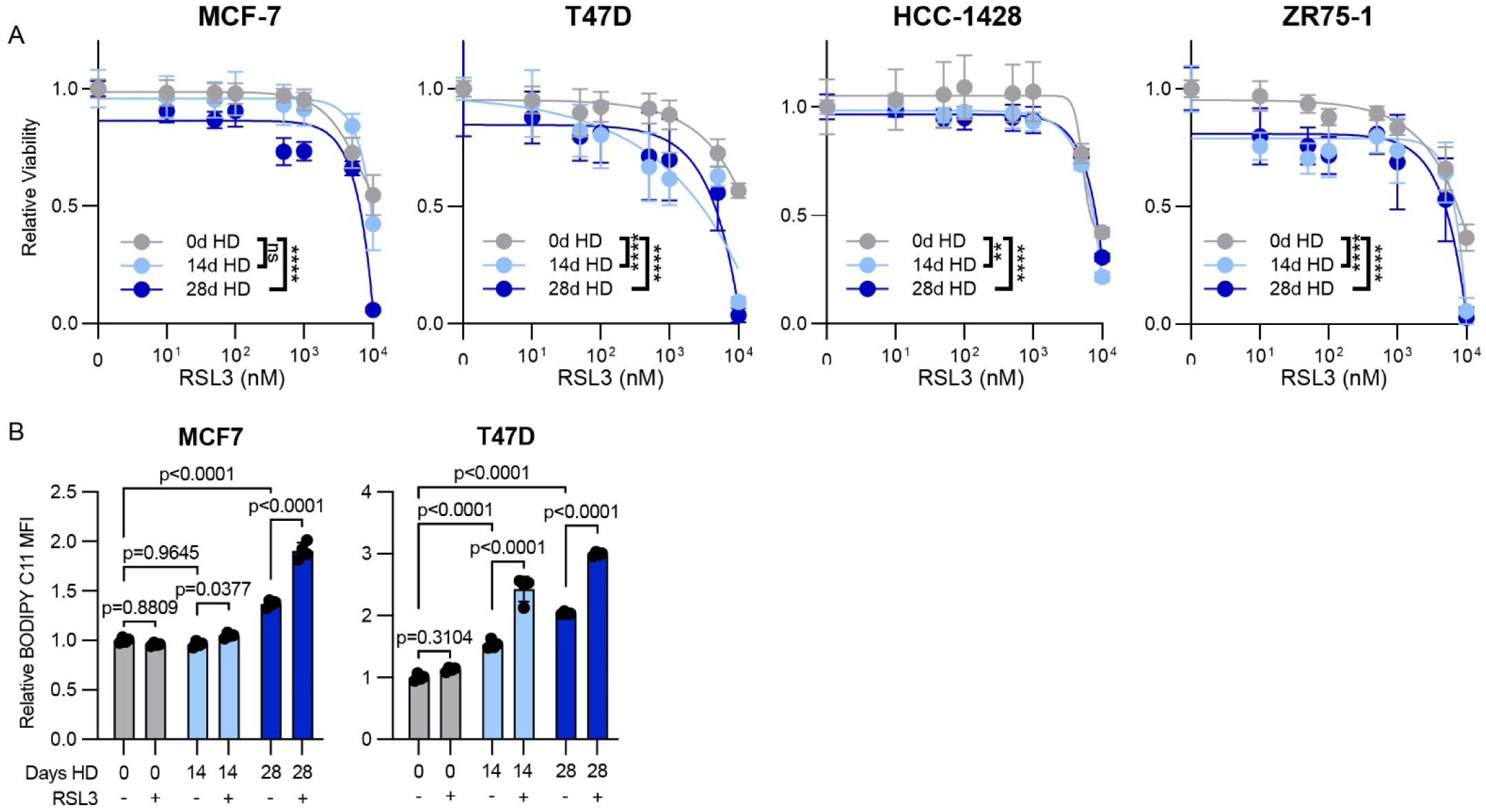
HD persister cells are modestly sensitized to RSL3. A) Viability analysis of cells pre-HD for 0-28 d, before reseeding and treatment with a dose range of RSL3 for 3 d. Viability was assessed by SRB staining. Data represent mean of sextuplicates + SD. Areas under the curves were compared by Bonferroni-adjusted posthoc test. B) Flow cytometry analysis of cells pre-HD for 0-28 d before treatment with 10 μM RSL3 for 4 h. Cells were then stained for BODIPY C11 and assayed by flow cytometry.

**Supplemental Figure 5:**
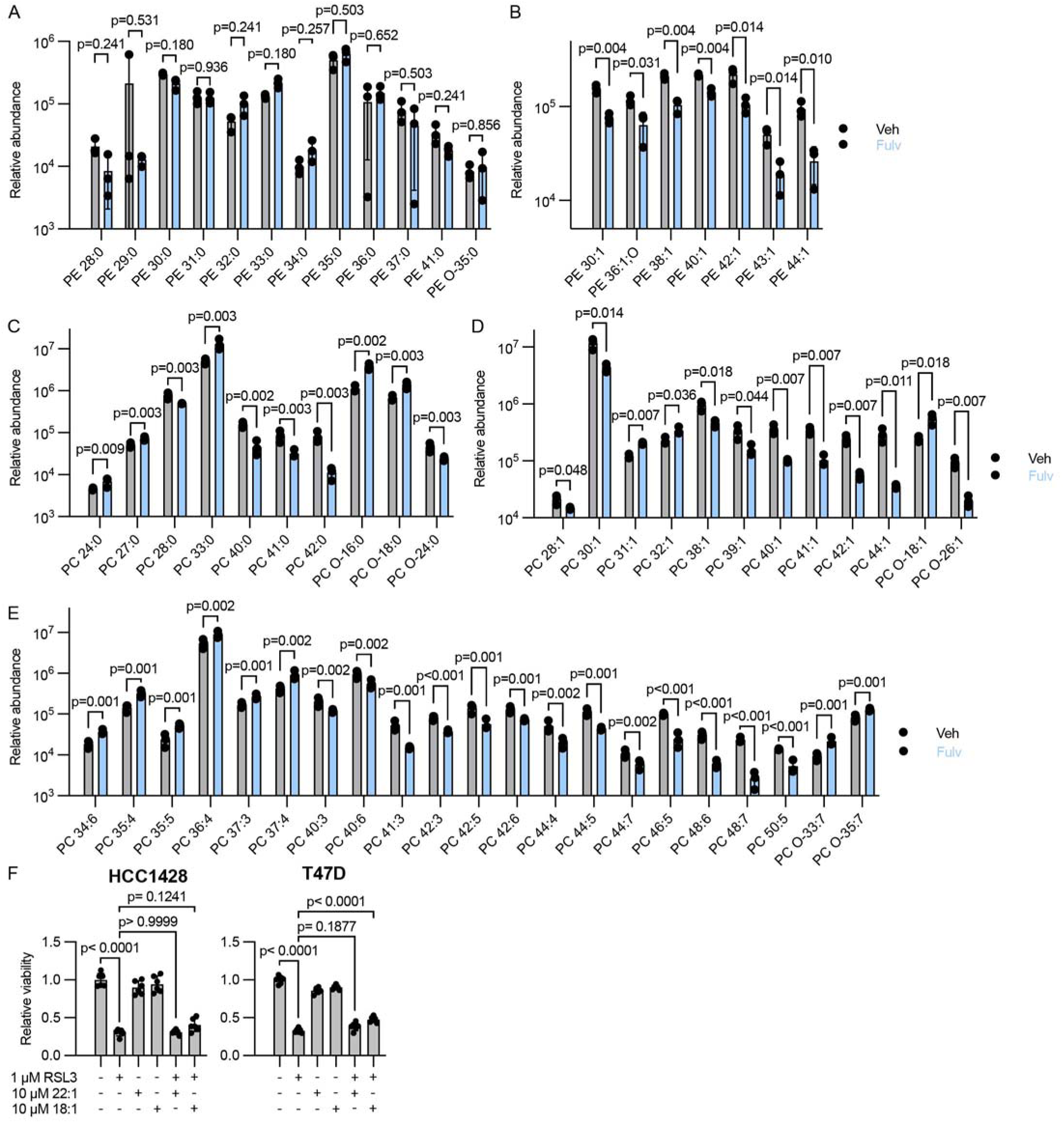
Endocrine therapy induces phospholipid remodeling including loss of PE and PC phospholipid species less labile to peroxidation. A-D) Relative abundance of PE SFA (A), PE MUFA (B), PC SFA (C), PC MUFA (D), and PC PUFA (E) from Figure 3.8A. p-values were calculated by multiple two-sided t-test with 5% FDR. F) Relative viability of HCC1428 and T47D cells pre-treated with fulvestrant for 7 d prior to subsequent treatment +/− RSL3, erucic acid (22:1), and oleic acid (18:1).

**Supplemental Figure 6:**
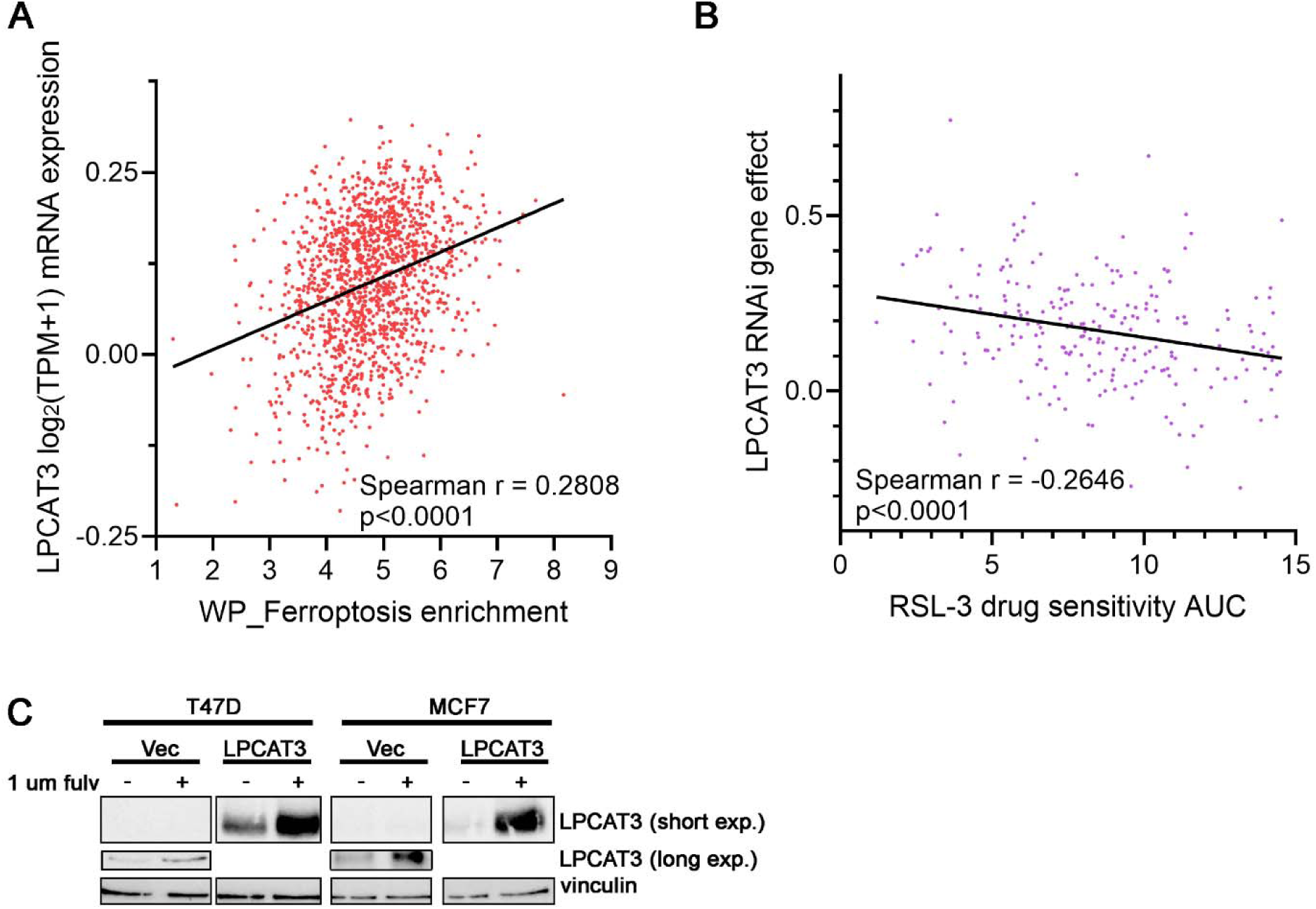
LPCAT3 overexpression induces phospholipid remodeling. A) *LPCAT3* mRNA expression levels across 1,516 cancer cell lines were compared to gene set enrichment scores for the 64-gene WP_FERROPTOSIS signature within the DepMap Portal. B) *LPCAT3* gene effect values (derived from sernsitivity to RNAi targeting *LPCAT3* in Achilles+DRIVE+Marcotte, DEMETER2) across 235 cancer cell lines were compared to area under the curve (AUC) values in response to a dose range of RSL3 (1S,3R-RSL-3; CTRP:609060; in Cancer Target Discovery and Development (CTD^2^) Network). Lower AUC indicates increased sensitivity to RSL3. C) Cells stably overexpressing LPCAT3 (or vector control) were treated ± 1 uM fulv for 2 d. Lysates were analyzed by immunoblot.

**Supplemental Figure 7:**
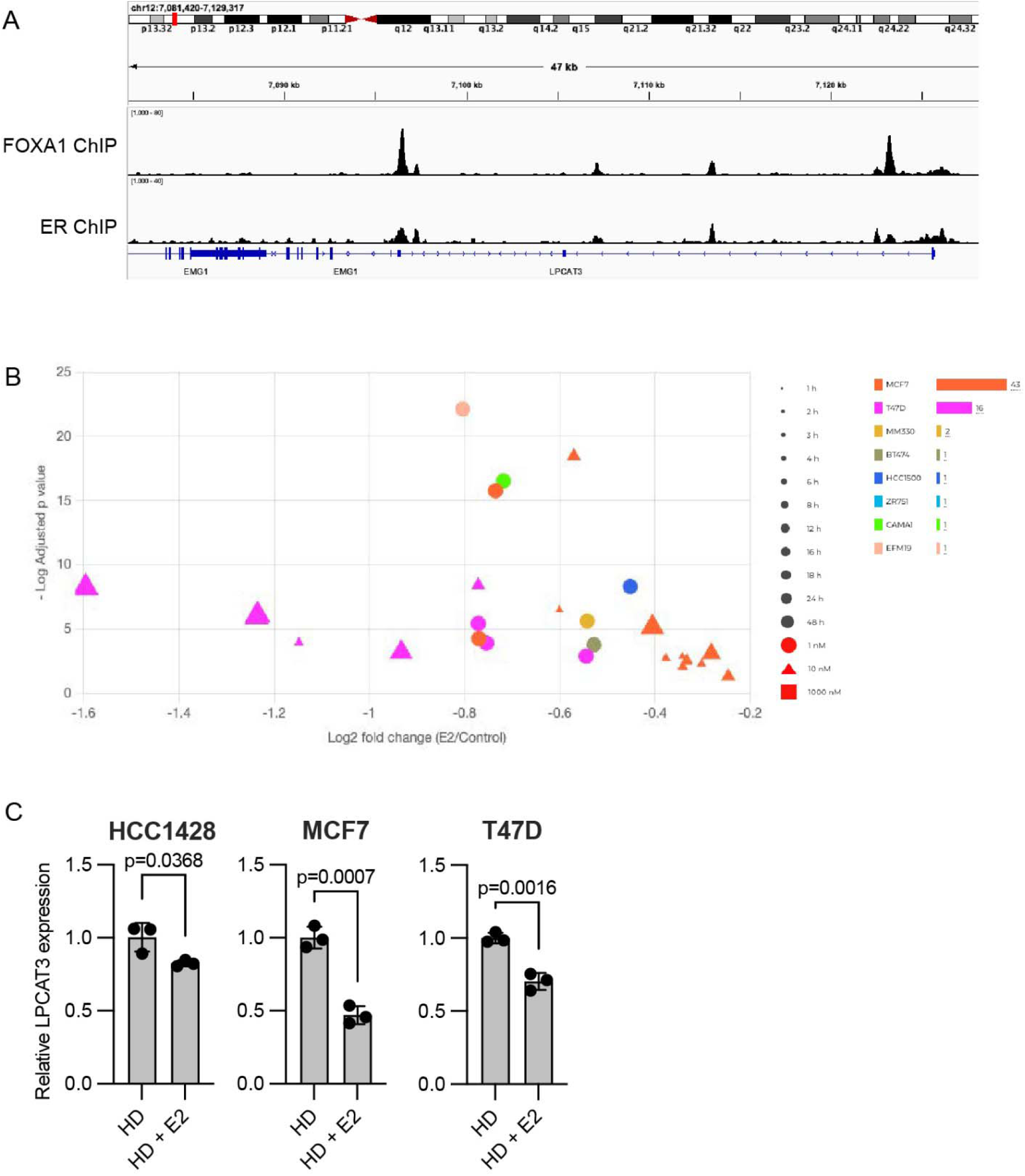
ER signaling regulates LPCAT3 expression. A) Plot of ChIP-seq data downloaded from GSE72249 displaying peaks around the transcription start site of *LPCAT3* in T47D cells following treatment with E2. B) Volcano plot of collective *LPCAT3* expression from RNA-seq datasets as analyzed using the Estrogene database. C) RT-qPCR analysis of RNA extracted from cells treated in with hormone deprivation for 7 d followed by ± 1 nM E2 for 2 d. Data are shown as mean of triplicates ± SD.

**Supplemental Figure 8:**
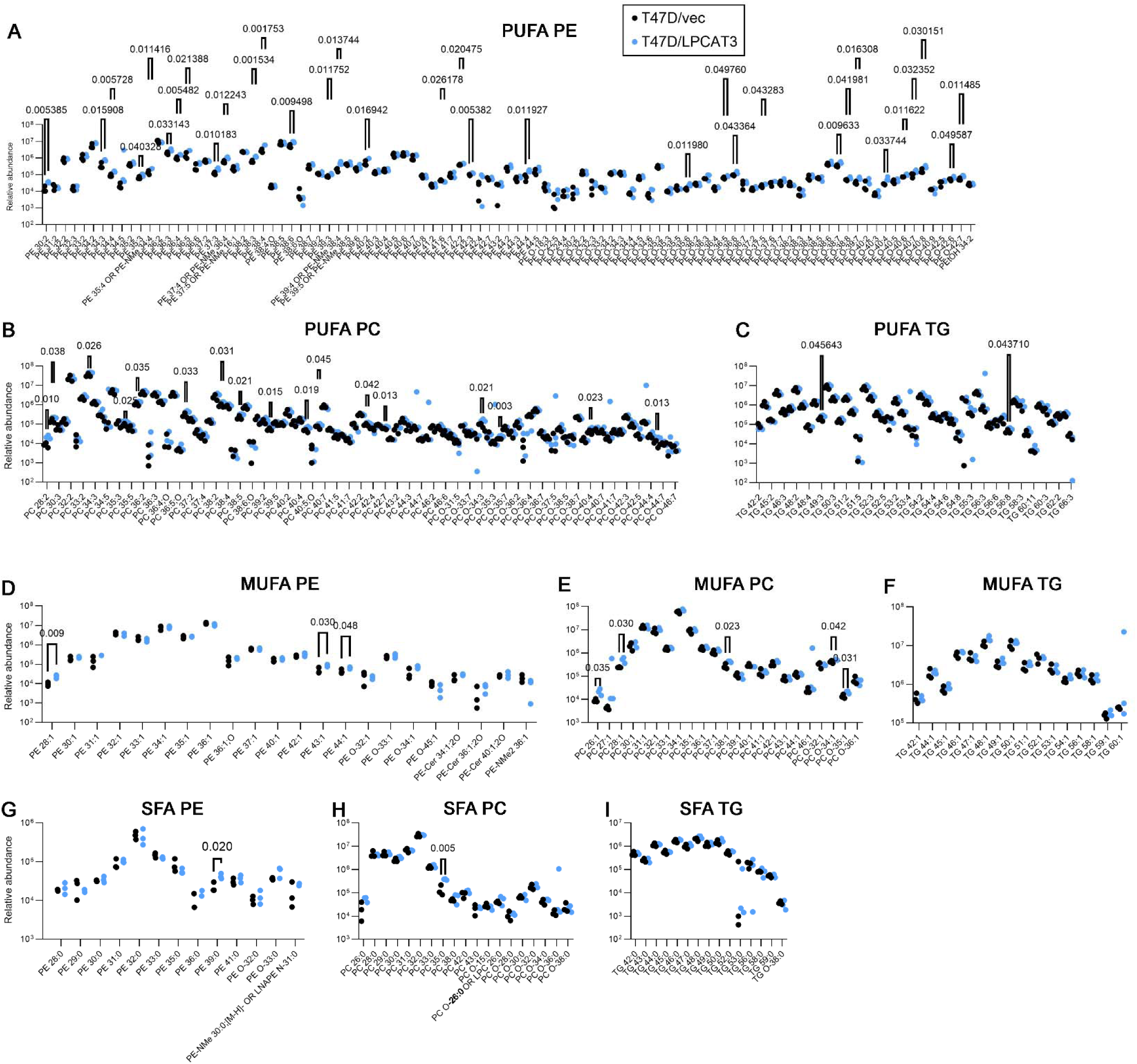
LPCAT3 overexpression induces phospholipid remodeling. Lipids were extracted from untreated T47D/vec and T47D/LPCAT3 cells. Shown are relative abundances of PE PUFA (A), PC PUFA (B), TG PUFA (C), PE MUFA (D), PC MUFA (E), TG MUFA (F), PE SFA (G), PC SFA (H), and TG SFA (I). p-values were calculated by multiple two-sided t-test with 5% FDR. Only comparisons with p≤0.05 are noted.

**Supplemental Figure 9:**
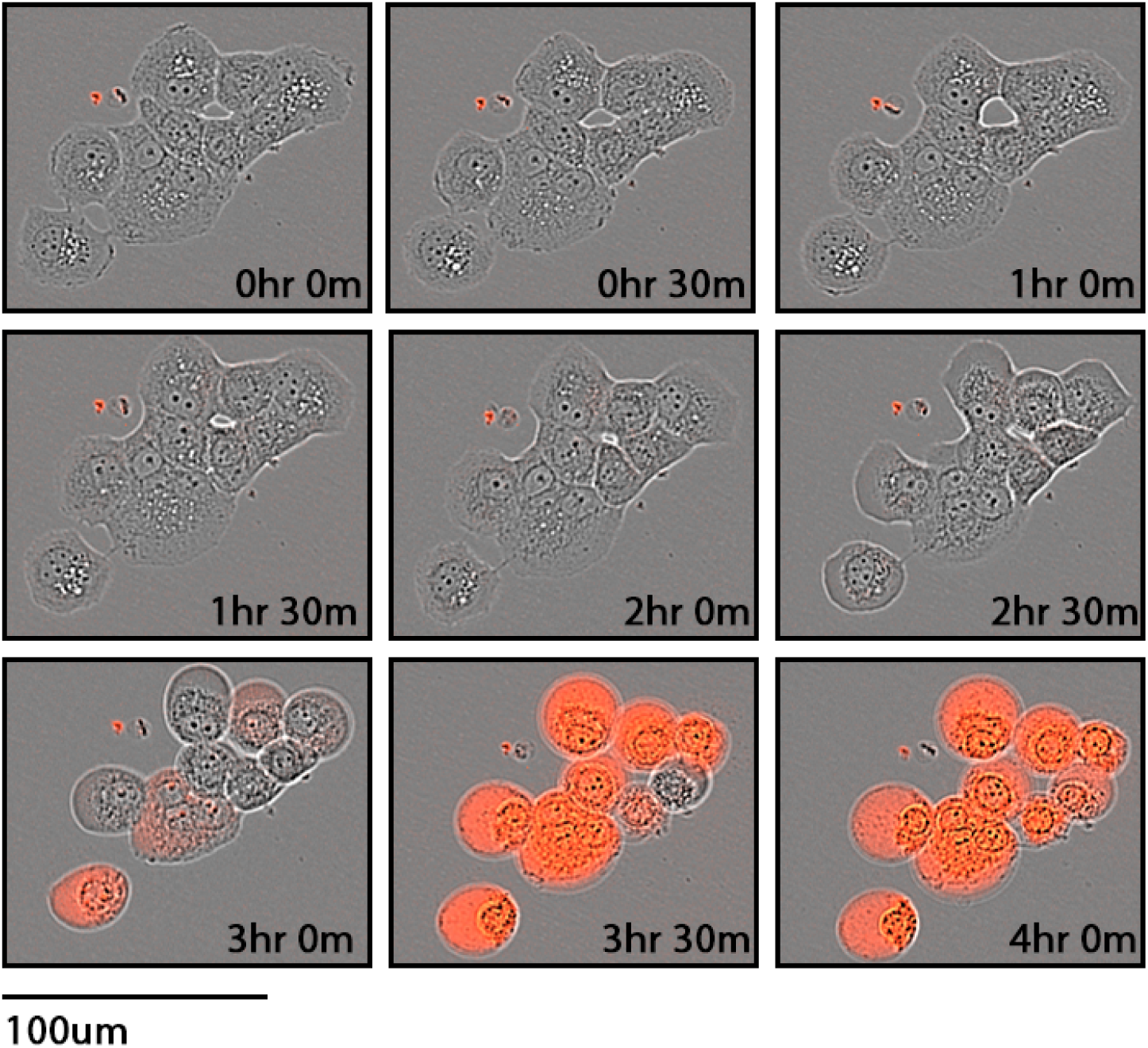
RSL3 induces ferroptosis within hours. T47D/LPCAT3 cells were pretreated with fulv for 3 d, and then incubated with Cytotox dye and simultaneously co-treated with 1 uM RSL3. Dye uptake, which is indicative of plasma membrane rupture, was detected by Incucyte serial imaging every 30 min. Representative images of cells with eventual dye uptake are shown.

**Supplemental Figure 10:**
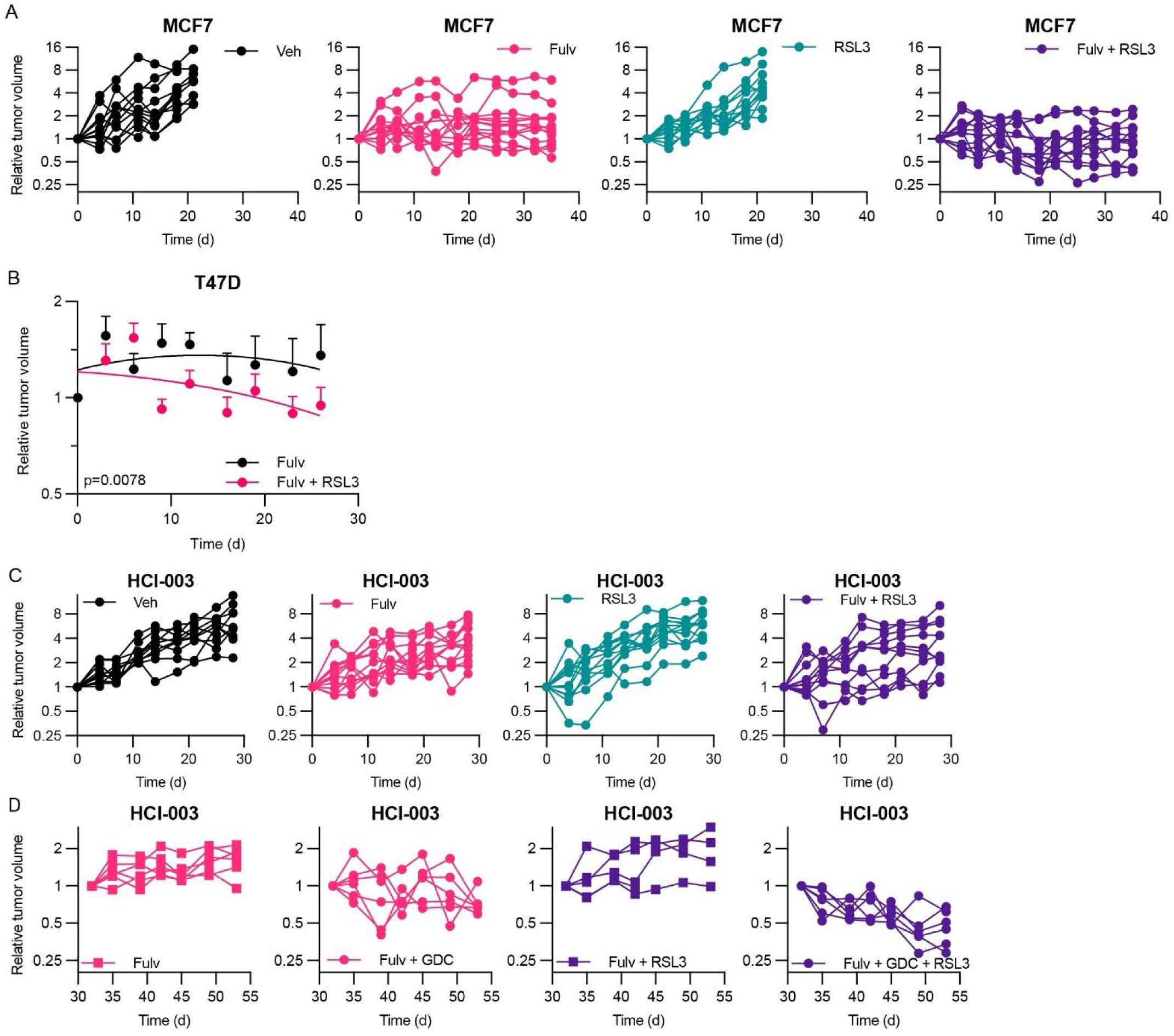
Growth curves of xenografts treated with fulvestrant and RSL3. (A) Individual tumor volume curves of MCF7 xenografts treated with fulvestrant QW and RSL3 QDx5. (B) Plot of tumor volume curves of T47D xenografts treated with fulvestrant QW ± RSL3 QDx5. Data are shown as mean + SEM (n = 10/group). Slopes determined by linear regression of log-transformed data were compared by t-test. (C) Individual tumor volume curves of HCI-003 PDXs treated with fulvestrant QW and RSL3 QDx5. (D) Individual tumor volume curves of HCI-003 PDXs treated with fulvestrant QW, GDC-0941 QDx5, and RSL3 QDx5.

## Notes

### Competing Interest Statement

The authors have declared no competing interest.

